# The role of ultradian rhythms in post-deprivation rebounds and diurnal rhythms of sleep and wakefulness in rats

**DOI:** 10.1101/2024.02.04.578825

**Authors:** Joonbum Lim, Richard Stephenson

**Author notes:** Corresponding author E-mail address (Richard Stephenson).

## Abstract

The temporal organization of ultradian rhythms in sleep and wakefulness during post-sleep deprivation (TSD) rebound were investigated in 15 rats under contant bright light (LL). Following baseline recordings, rats were subjected to TSD using gentle manual stimulation. Post-TSD rebounds in cumulative wakefulness (WAKE), rapid eye movement sleep (REM) and non-REM sleep (NREM) were analyzed in WAKE-dominant (υ_w_) and sleep-dominant (υ_s_) ultradian phases. Rebounds in WAKE and NREM were present only when data were analyzed on a full ultradian cycle basis, and were absent in υ_s_ and υ_w_ phases alone. These rebounds were approximately 50% complete and not proportional to TSD excess/deficit. Rebounds in REM were present in full ultradian cycles and partially expressed in υ_s_ but absent in υ_w_. REM rebounds fully compensated for REM deficit. Rebounds were mediated mainly by a reduction in the duration of the υ_w_ ultradian phase, and by decreased probability of arousal in the υ_s_ ultradian phase. These mechanisms were also found to partially mediate diurnal rhythms in 10 rats under a 12:12 h LD cycle.

This study implicates an ultradian timing mechanism in the control of post-TSD rebounds and suggests that rebounds in all three states are mainly mediated by post-TSD adjustments in WAKE-promoting mechanisms. Ultradian rhythms should be taken into account to avoid errors in data analysis.

**Highlights:**

- Sleep-wake state exhibits circadian rhythms and ultradian rhythms.
- These rhythms interact with rebounds after sleep deprivation.
- Circadian amplitude and sleep rebound are partially mediated by ultradian timing.
- Arousal-related processes control these sleep-wake patterns in both states.
- Measuring ultradian rhythms is necessary for accurate analysis of data.

## 1. Introduction

Sleep-wake states (wakefulness, WAKE, rapid-eye movement sleep, REM and non-REM sleep, NREM) alternate over time in a pattern that exhibits probabilistic and deterministic characteristics. For example, in the short-term, the exact timing of state transitions is not predictable, whereas over longer time intervals periodic diurnal rhythms, quasi-periodic ultradian rhythms, and monotonic “rebound” responses following acute total sleep deprivation (TSD) have all been well described in mammals[1-7]. These quantitative features of sleep-wake patterns are expressed at least partly independently but they generally occur concurrently. The co-dependence of circadian rhythms and rebound responses is well established and the majority of studies explicitly account for this in their design and interpretation [7-12]. Recent studies have emphasized the significance of ultradian rhythmicity in state expression[6, 13-21], but the interactions between ultradian rhythms and other components of the sleep-wake pattern have not yet been examined rigorously. For example, interactions between ultradian rhythms and post-TSD rebounds remain to be investigated, and were the focus of the present study.

Whilst it is often claimed that sleep duration is regulated in proportion to prior wakefulness[22], there is little evidence for a positive serial correlation between WAKE bout durations and subsequent sleep episode durations in rodents[9, 23, 24], a conclusion supported in the present study. This implies that “prior wakefulness” is not simply the duration of the immediately preceding WAKE bout, and suggests that post-TSD rebounds are mediated by an as yet poorly characterized mechanism that operates over a longer (multi-bout) timescale. We hypothesize that the ultradian rhythm participates in this longer-range process.

We have previously postulated a probabilistic toy model[19] for the origin of quasi-periodicity in sleep-wake ultradian rhythms in which intervals of predominant wakefulness (the υ_w_ ultradian phase) alternate with intervals characterized by a predominance of sleep (the ultradian υ_s_ phase). We speculated that the υ_s_ phase might equate to the putative sleep homeostat. This model, which is analogous to the two-process model[25], predicts that enforced wakefulness will be followed by a rebound in which the ultradian υ_s_ phase is prolonged.

All of the above-mentioned long-term patterns of sleep-wake expression emerge from sequences of individual bouts of WAKE, NREM and REM that are subject to short-term control of state maintenance and inter-state transition. The timing of state transitions appears to be controlled, at least in part, by one or more stochastic processes[15, 20, 26-36] and the longer term patterns can therefore be understood and quantified in terms of controlled variations in transition probabilities. We have previously emphasized the importance of the non-linear relation between transition probability and state bout duration for the relative influence of the various transition events in overall state control[37]. We concluded that the probability of state maintenance of the “long” WAKE sub-type has the greatest potential impact in the control of all three states in relation to circadian rhythms and post-TSD rebound responses, while post-NREM transition trajectory (quantified as the normalized probability of post-NREM arousal, 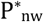) is potentially important in the control of REM expression.

Circadian rhythms have a profound influence on state transition probabilities and complicate the analysis of long-term rebound responses, so in this study we recorded from rats maintained under constant bright light (LL) to suppress circadian rhythms. Our objective was to determine whether a rebound response would occur under these conditions, and if so, to test the above-mentioned model predictions. Contrary to expectations, adjustments in state bout durations and frequency within each ultradian phase were found to have a surprisingly small impact on long-term expression of post-TSD responses under these experimental conditions. Furthermore, also contrary to model predictions, the duration of the wake-dominant (υ_w_) phase played an important causal role in the expression of rebounds of all three vigilance states. This study suggests that ultradian timing of wakefulness is an important variable in the control of post-TSD rebounds of both wakefulness and sleep.

## 2. Methods

All experimental procedures were performed in strict accordance with the guidelines established by the Canadian Council on Animal Care and were approved by the animal care committee at the University of Toronto.

### 2.1. Animal maintenance and experimental protocols

Eighteen rats were used, but 3 were rejected due to substandard signal quality or equipment failure. Thus, data from a total of 15 male Sprague-Dawley rats were analyzed in this study. Rats were adults, ranging from 24 to 40 weeks old at start of study, with relatively stable body mass throughout; mean ±S.E.M. 713 ±21 g at start of recordings and 709 ± 18 g at end of study. They were kept individually in standard laboratory cages with constant environmental conditions free of time cues, and had unrestricted access to chow and water throughout the study. Cages were housed in a sound-attenuated room at 23 ±1 **°**C and, except during acute total sleep deprivation (TSD), rats experienced minimum (irregular) disturbance for technical procedures. At least 10 weeks before experiments began, and continuing uninterrupted until completion of recordings, rats were maintained in constant bright light (LL, approximately 650 lux at cage floor).

Animals were recorded continuously for at least 9 days (initial baseline, BL) before being subjected to acute sleep deprivation (TSD1), followed by 7 days undisturbed (REC1), followed by a second, similar sleep deprivation (TSD2), followed by a final 7 days undisturbed (REC2).

The 15 rats were divided into 3 groups of 5 animals each: SD2, SD4 and SD6. Animals in each group were housed in a single environmental room and were studied simultaneously. Sleep deprivation was achieved using gentle manual stimulation as follows. Electroencephalogram (EEG) and electromyogram (EMG) recordings were displayed in real time, together with direct visual observation of animal behaviour. Sleep onset (drowsiness) was defined as any time when a rat ceased activities and assumed a relaxed posture, EMG activity decreased noticeably, and EEG amplitude and slow wave (< 6 Hz) activity began to increase for two or more epochs (i.e. 10 s). At this point, wakefulness was enforced using the minimum effective stimulus, beginning with touching the whiskers with a soft brush and progressing if necessary to pushing the fore- or hindlimbs with the handle of the brush. Sometimes, movement of the researcher within the environmental chamber was sufficient to alert one or more animals, and this precluded the measurement of stimulus frequency and intensity. This protocol was sustained for 2 h (SD2 group), 4 h (SD4 group) or 6 h (SD6 group).

### 2.2. Instrumentation

All animals were instrumented, under isoflurane anaesthesia, with fully implanted biotelemetry devices (model TL10M3-F50-EET, Data Science International, St. Paul, MN) for acquisition of EEG (fronto-parietal bipolar electrodes; coordinates, expressed relative to bregma: AP +2mm / ML 2mm, AP - 4mm / ML 2mm), EMG (nuchal bipolar electrodes) and intraperitoneal temperature (T_b_). The surgical procedures were exactly as described in detail previously[38]. During post-operative recovery, animals were administered analgesic (ketoprofen, 3mg/kg, sc) twice per day for the first two days.

### 2.3. Data acquisition

Data were acquired (sampling frequency, 400 Hz) and conditioned in 5 s epochs using custom software (LabView version 7, National Instruments Corp., Austin, TX). Data processing and sleep scoring were performed as described previously[10, 39]. Briefly, offline automated sleep scoring was conducted using our rat sleep autoscoring algorithm written in spreadsheet format (ratSAS; Microsoft Excel, v. 2011, Microsoft Corp., Redmond, WA, USA). This algorithm has been described in detail and validated for use under the conditions of this study[39].

Ten days after surgery the telemetry device was activated via a magnetic switch, and a brief (∼3h) preliminary recording taken to verify data quality. Visual analysis of the preliminary raw data was performed to check the accuracy of the automated scoring system in each rat and all animals used in the study had autoscore versus visual rater concordance >90%. In both automated and visual scoring each epoch was assigned to one of four categories; wakefulness (WAKE), rapid eye movement sleep (REM), non-REM sleep (NREM), and artifact (ART). Artifacts were identified algorithmically using the method described previously [10], which is based on Tukey’s non-parametric outlier detection protocol[40]. All epochs identified as ART (1.1 ± 0.3 % of all BL epochs recorded) were subsequently reclassified by direct visual inspection based on preceding and subsequent EEG and EMG patterns and 99.6 ± 0.2 % of identified ART were judged to have occurred during wakefulness. The behavioural scores of reclassified ART epochs were included in analyses of behaviour, but the electrophysiological variables were not.

In this study, a “bout” of each state is defined as any uninterrupted interval (one or more epochs) in a given state of WAKE, NREM or REM. Uninterrupted intervals of sleep consisting of more than one bout (i.e. sequences of NREM and REM) are referred to as “sleep episodes”.

### 2.4. Data Analysis

All statistical tests were performed using either Microsoft Excel spreadsheets or GraphPad Prism statistical software (version 6.0f for mac OSX, GraphPad Software, San Diego, CA, USA). Group data were considered to be normally distributed when they “passed” (i.e. P > 0.05) at least one of the following tests; D’Agostino and Pearson omnibus K2 test, Shapiro-Wilk test and Kolmogorov-Smirnov test. This result was used to justify the use of parametric or non-parametric statistical tests and summary statistics.

#### 2.4.1 Circadian and ultradian periodicities in sleep-wake state and body temperature time series

In previous studies we concluded that ultradian rhythms are quasi-periodic[6]. Commonly used techniques such as maximum entropy spectral analysis, autocorrelation, chi-square periodogram and Lomb-Scargle periodogram are sub-optimal when data are non-stationary[41-43], because they cannot distinguish between concurrent or alternating periodicities when multiple peaks are present in the periodogram. For this reason we used continuous wavelet analysis in the present study (R-project, WaveletComp package[44], implemented using RStudio version 2021.9.1.372 (RStudio, PBC, Boston, MA)). This method enabled us to quantify the evolution of spectral content over time across the study. In this part of the study, the time series were collapsed into 10 min time bins[45]. The fractional content of WAKE (F_WAKE_ = 1-F_SLEEP_) and mean T_b_ data in each 10 min time bin were analyzed using the *analyze*.*wavelet* routine (see [44] for details) with spectral resolution dj = 1/20 per octave, over the period range 1 – 32 h. In addition to the time-varying wavelet power spectrum, global wavelet spectral density (and associated statistical probability) was obtained as the time-averaged wavelet power spectrum. The latter was used to estimate period of circadian rhythms during BL and during days 3-7 of REC1 and REC2. Cross-wavelet power spectra were also obtained using *analyze*.*coherency* to evaluate joint spectral power and phase lags between F_WAKE_ and mean T_b_.

#### 2.4.2. Circadian phase in the sleep-wake time series

Circadian rhythms were not readily discernable by visual inspection of the F_WAKE_ and T_b_ time series. However, statistically significant low amplitude periodicities within the circadian range (18 – 32 h) were revealed in some parts of the F_WAKE_ and/or T_b_ wavelet power spectra. Average circadian period was identified for each animal in the corresponding global wavelet power spectra and this was used in least-squares regression of F_WAKE_ data (after zero-reset by subtraction of the time series mean) using the method described by Lomb[46]. The time of descending zero-cross of the fitted F_WAKE_ sine wave was assumed to be circadian time (CT) 0 h, approximating time of lights on in rats held in a 12:12 h LD cycle. The time of ascending zero-cross of the fitted F_WAKE_ sine wave was assumed to be circadian time CT 12 h, approximating dark onset in rats held in a 12:12 h LD cycle (1 circadian h = period / 24). Data were then designated as subjective L and subjective D across the study. Circadian phase was noted at the start of TSD and the start of REC.

In order to estimate the extent of suppression of rhythms by LL, we compared the data with previously published data[37] from 10 rats kept under 12:12 h LD cycle. All technical protocols used in the LD studies were closely similar to those used in the present study. Data from 7 full BL days were subjected to an identical analysis to compare high amplitude entrained diurnal rhythms under LD with strongly attenuated free-running circadian rhythms under LL. Emphasis here was on the relation between ultradian and circadian characteristics under the two conditions.

#### 2.4.3. Ultradian phase in the sleep-wake time series

A binary score was used for ultradian cycles and the following procedure was used to mark the transitions between υ_w_ and υ_s_ ultradian phases in the F_WAKE_ time series over the entire study. First of all, the time series was smoothed using a 4 h centered moving average to extract circadian periodicities. This low-pass average was subtracted from the original time series to reset the mean to zero and leave only the residual ultradian periods (i.e. < 8 h). A scaling factor ([1-mean]/mean) was then applied to all sub-zero values in order to balance the magnitudes of peaks and troughs of the ultradian F_WAKE_ waveform. This was needed in order to avoid spurious lags in transition timing in the subsequent stage of this scoring protocol. Phase transitions were next defined by zero-crossing of a 90 min centered moving average through the rescaled zero-reset F_WAKE_ time series. Supra-zero intervals were scored as υ_w_ and sub-zero intervals were scored as υ_s_. To avoid “dithering” (i.e. erroneous rapid phase transitions) during parts of the time series where moving average F_WAKE_ remained close to zero, a minimum phase duration criterion of 40 min was applied. Note that rescaled zero-reset data were used solely for the purpose of categorical ultradian binary phase scoring and were not used in any other calculations in this study.

Ultradian period, and the durations of the υ_s_ and υ_w_ phases were recorded for each one of 12 full ultradian cycles before and after TSD. The proportion of each ultradian υ_s_-υ_w_ cycle consisting of the υ_w_ phase (duty cycle, D_c_) was also recorded.

#### 2.4.4. Acute TSD

Total sleep deprivation (TSD) was enforced on a per group basis. That is, all the rats in a given group were deprived of sleep simultaneously. Since the rats in each group were free-running, the TSD protocol was initiated in each animal at random circadian and ultradian phases. All rats were awake for a variable amount of time before the start of the TSD protocol, either spontaneously or due to disturbance associated with entry of the researcher into the room. This “pre-stimulus wakefulness” was considered to be part of the TSD interval. At the end of the designated TSD stimulus interval, rats were left to sleep undisturbed. In all cases, they remained awake for a variable amount of time before sleep onset and this interval between the end of stimuli and the onset of sleep was also included in calculations of WAKE excess and sleep deficit.

TSD-induced WAKE excess and sleep deficit were calculated as follows. In each animal, rates of expression (i.e. fractional time in state) of WAKE (F_WAKE_), NREM (F_NREM_) and REM (F_REM_) were calculated for full υ_s_-υ_w_ ultradian cycles in the BL interval, and averaged across cycles, with care taken to match the circadian phases of the BL samples with those of the TSD interval. Predicted time in state was then calculated as: BL mean fractional time in state x TSD interval / ultradian cycle duration. Overall excess/deficit was calculated for each state by subtraction of the predicted time in state from the observed time in state.

The experimental stimuli used to enforce TSD strongly masked both sleep-WAKE behaviour and T_b_, thereby precluding a direct measure of ultradian phase during the TSD interval. To obtain predicted values and corresponding excess/deficit estimates for each of the ultradian phases separately, we assumed that the ultradian cycle continued unchanged from BL during TSD. The overall TSD WAKE excess and sleep deficits calculated above were partitioned between fictive ultradian phases as follows. Using circadian phase-matched BL data, we recorded ultradian duty cycle (D_c_ = υ_w_ duration / υ_s_-υ_w_ cycle duration), and mean ultradian phase-specific F_WAKE_, F_NREM_ and F_REM_. Duty cycles were used to divide total TSD interval into predicted time in each ultradian phase. These predicted times in phase were then multiplied by predicted phase-specific F_WAKE_, F_NREM_ and F_REM_ values to obtain total predicted time in each state within each fictive ultradian phase across the TSD interval. Next, the *observed* F_WAKE_, F_NREM_ and F_REM_ values during TSD were multiplied by *predicted* time in each ultradian phase to obtain an estimate of total time in each state within each fictive ultradian phase across the TSD interval. Ultradian phase-specific excess/deficits were then taken as the observed – predicted times in state for each fictive phase.

EEG “slow wave activity” during NREM sleep (SWA_NREM_) is widely assumed to be an index of NREM intensity[47]. According to this hypothesis, total NREM sleep deficit during TSD is the difference between cumulative SWA_NREM_ expressed during the TSD interval and the corresponding amount predicted by extrapolation of BL levels over the same interval. SWA_NREM_ was estimated as delta amplitude (μVrms of the EEG frequency band 1.5-6 Hz[48]) during NREM epochs. Cumulative SWA_NREM_, referred to here as NREM “slow wave energy” (SWE_NREM_) was recorded as the sum of SWA_NREM_ in NREM epochs over the TSD interval. Predicted SWE_NREM_ during TSD was calculated as TSD duration x predicted F_NREM_ x predicted SWA_NREM_. Total SWE_NREM_ deficit was the difference between observed and predicted values over the TSD interval.

#### 2.4.5. Conventional analysis of rebound

Fractional times in state (F_WAKE_, F_NREM_, F_REM_) were calculated in non-overlapping 2 h time bins over the interval 72 h before TSD (BL) to 72 h after TSD (REC). Following convention, we assumed that post-TSD rebound was the difference between observed state expression and that predicted by extrapolation of mean BL rate of state expression over the same time interval, taking circadian phase into account. Thus, for each circadian phase, mean F_WAKE_, F_NREM_ and F_REM_ were calculated over the 36 BL time windows and subtracted from respective F_WAKE_, F_NREM_ and F_REM_ observed in each of the circadian phase-matched REC windows. The differences obtained were cumulated over REC time and subtracted from TSD-induced WAKE excess, NREM deficit and REM deficit (see *2*.*4*.*4. Acute TSD*, above) to yield cumulative rebounds. Data are expressed as difference from BL.

#### 2.4.6. Ultradian phase-adjusted analysis of rebound

This was similar to the conventional analysis described in section *2*.*4*.*5*. above, but in this case ultradian phase was taken into account when calculating the predicted values during REC. In BL data, mean F_WAKE_, F_NREM_ and F_REM_ were derived separately for υ_s_ and υ_w_ ultradian phases, in each of the subjective L and D circadian phases. For each of the 36 time windows during REC, the fraction of time taken up by each ultradian phase was recorded, then predicted F_WAKE_, F_NREM_ and F_REM_ were calculated for each window as (fraction time υ_s_ x mean BL F_i_υ_s_)+(fraction time υ_w_ x mean BL F_i_υ_w_) for each state i. Data are expressed as observed – predicted difference.

#### 2.4.7. Ultradian phase-specific analysis of rebound

The results of the analyses described above demonstrated that ultradian rhythms are implicated in post-TSD rebounds. We therefore undertook a more detailed investigation of the role of ultradian phase as follows.

Calculations of rebounds were repeated on an ultradian phase-by-phase and cycle-by-cycle basis. Thus, 12 complete υ_s_ and υ_w_ phases were analyzed immediately before and after TSD. Phase durations, and fractional time in state were calculated for each individual ultradian phase interval and for corresponding complete υ_s_-υ_w_ ultradian cycles. The duty cycle (D_c_) was calculated for each BL and REC ultradian cycle. Mean values of fractional time in state were calculated over the BL cycles, and used as predicted values during REC, taking circadian phase into account. Thus, during REC, predicted time in state in a given ultradian phase was the product of phase duration x BL mean fractional time in state for the same phase, again matching for subjective circadian phase. As before, rebounds were then calculated and expressed as observed – predicted values.

Durations (min) and frequency (bouts/h) of bouts of WAKE, NREM and REM were recorded for each of the 12 υ_s_ and υ_w_ phases, and the corresponding full υ_s_-υ_w_ cycles, immediately before (BL) and after (REC) TSD. Values during REC were expressed as difference from phase-specific mean BL values.

Normalized probability of arousal 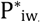= n_iw_/n_i_, where index w denotes WAKE and index i = n for NREM, and i = r for REM) was estimated as the fractions of NREM bouts and REM bouts that were followed by WAKE. Thus, 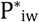, where n_iw_ is number of bouts of state i that were followed by WAKE and n_i_ is the total number of bouts of state i. 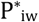 is the additive inverse of the normalized probability of sleep cycling (i.e. NREM to REM transitions, 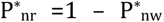, and REM to NREM transitions, 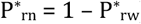.

Serial correlations were computed (Spearman rank correlation coefficient, ρ) to quantify statistical associations between WAKE bout durations and the succeeding sleep episode duration in each ultradian phase. In addition, Spearman rank correlation coefficient was calculated for REM bout duration versus succeeding inter-REM interval (IRI, comprising one or more bouts of WAKE and/or NREM). Due to small sample sizes within each individual ultradian phase, we pooled BL cycles, as well as the first 6 REC ultradian cycles encompassing approximately the first circadian day of rebound.

SWA_NREM_ was quantified on an ultradian phase-specific basis over the course of BL and REC. That is, SWE_NREM_ was calculated as the sum of SWA_NREM_ in NREM epochs in each ultradian phase. Predicted SWA_NREM_ during REC was assumed to be the circadian phase-matched mean BL SWA_NREM_ for each of υ_s_ and υ_w_. Predicted SWE_NREM_ during REC was calculated as ultradian phase duration x predicted F_NREM_ x predicted mean SWA_NREM_ in each phase. Rebound in SWE_NREM_ was calculated cumulatively over each sequential REC ultradian phase as SWE_NREM_ deficit – SWE_NREM_observed – SWE_NREM_predicted.

## 3. Results

### 3.1. Pre-TSD Baseline

#### 3.1.1.Baselines before first and second TSD procedures

Each rat was subjected to two similar TSD procedures separated by a 7-day recovery interval. Repeated measures ANOVA, with pre-planned post-hoc Holm-Sidak multiple comparisons tests were used to compare the two pre-TSD baselines. There were no statistically significant differences in any of the variables tested (ultradian period, durations of υ_w_ and υ_s_ phases, mean bout durations of WAKE, NREM and REM in each ultradian phase, mean bout frequency of WAKE, NREM and REM in each ultradian phase, mean F_WAKE_, F_NREM_ and F_REM_ in each ultradian phase and mean SWA_NREM_ in each ultradian phase). This confirms that recovery following TSD trial 1 was statistically complete more than 12 ultradian cycles before the beginning of TSD trial 2.

#### 3.1.2. Comparison of pre-TSD baselines between SD2, SD4 and SD6 groups

Ordinary one-way ANOVA, with pre-planned post-hoc Holm-Sidak multiple comparison tests were used to compare the three SD groups. There were no statistically significant differences between groups for ultradian period (F_(2,27)_ = 0.918, P = 0.41), υ_w_ phase duration (F_(2,27)_ = 0.052, P = 0.95) and υ_s_ phase duration (F_(2,27)_ = 2.741, P = 0.083).

Mean WAKE bout durations and mean REM bout durations did not differ significantly between SD2, SD4 and SD6 groups in υ_w_ (all P ≥ 0.36) and υ_s_ (all P ≥ 0.53). Mean NREM bout durations did not differ significantly between SD2, SD4 and SD6 groups in υ_w_ (F_(2,27)_ = 2.449, P = 0.11), but in υ_s_, mean NREM bout durations were 20% longer in the SD6 group than in the SD2 group (P = 0.014) whereas the SD4 group was statistically similar to both SD2 and SD6 (P = 0.41 and 0.07, respectively).

Mean WAKE bout frequency, mean NREM bout frequency and mean REM bout frequency did not differ significantly between SD2, SD4 and SD6 groups in υ_w_ (all P > 0.06) and υ_s_ (all P > 0.21). Mean F_WAKE_, F_NREM_ and F_REM_ did not differ significantly between SD2, SD4 and SD6 groups in υ_w_ (all P > 0.28) and υ_s_ (all P > 0.12).

Mean SWA_NREM_ did not differ significantly between groups in υ_w_ (F_(2,27)_ = 2.53, P = 0.098) and υ_s_ (F_(2,27)_ = 1.66, P = 0.21).

### 3.2. Wavelet analysis of wake-sleep and T_b_ time series

Fig 1A-C illustrates wavelet power spectra from a representative animal in LL. The figure demonstrates the variation in peak periods and peak power (ridges connecting spectral peaks over time are shown as black lines on a red background) over the course of a 25 day recording. The spectra show (i) that circadian rhythms were strongly suppressed, but not entirely abolished, in body temperature (T_b_, Fig 1A) and WAKE-sleep expression (F_WAKE_, Fig 1B). (ii) Spectral power of the circadian rhythm tended to wax and wane over time and was more pronounced overall in T_b_ than F_WAKE_. (iii) Global wavelet spectra (not shown) revealed that the free-running circadian period varied from 23.5 to 28.3 h among 15 animals. Mean free-running circadian period in LL was 26.1 ± 0.8 h and mean “circadian hour” was 65.2 ± 1.9 min duration. (iv) Non-stationary (quasi-periodic) ultradian oscillations were present throughout the study with peak periods varying considerably within the 2 – 8 h period range in each rat. (v) T_b_ and F_WAKE_ were strongly coherent over time, as demonstrated by the cross-wavelet power spectrum (Fig 1C), and the T_b_ waveform was slightly phase delayed relative to F_WAKE_ (indicated by the slight downward projection of the black arrows in Fig 1C). (vi) Spectral power tended to diminish over the first 12 – 24 h following TSD, suggesting a post-TSD change in ultradian waveform.

**Fig. 1.**
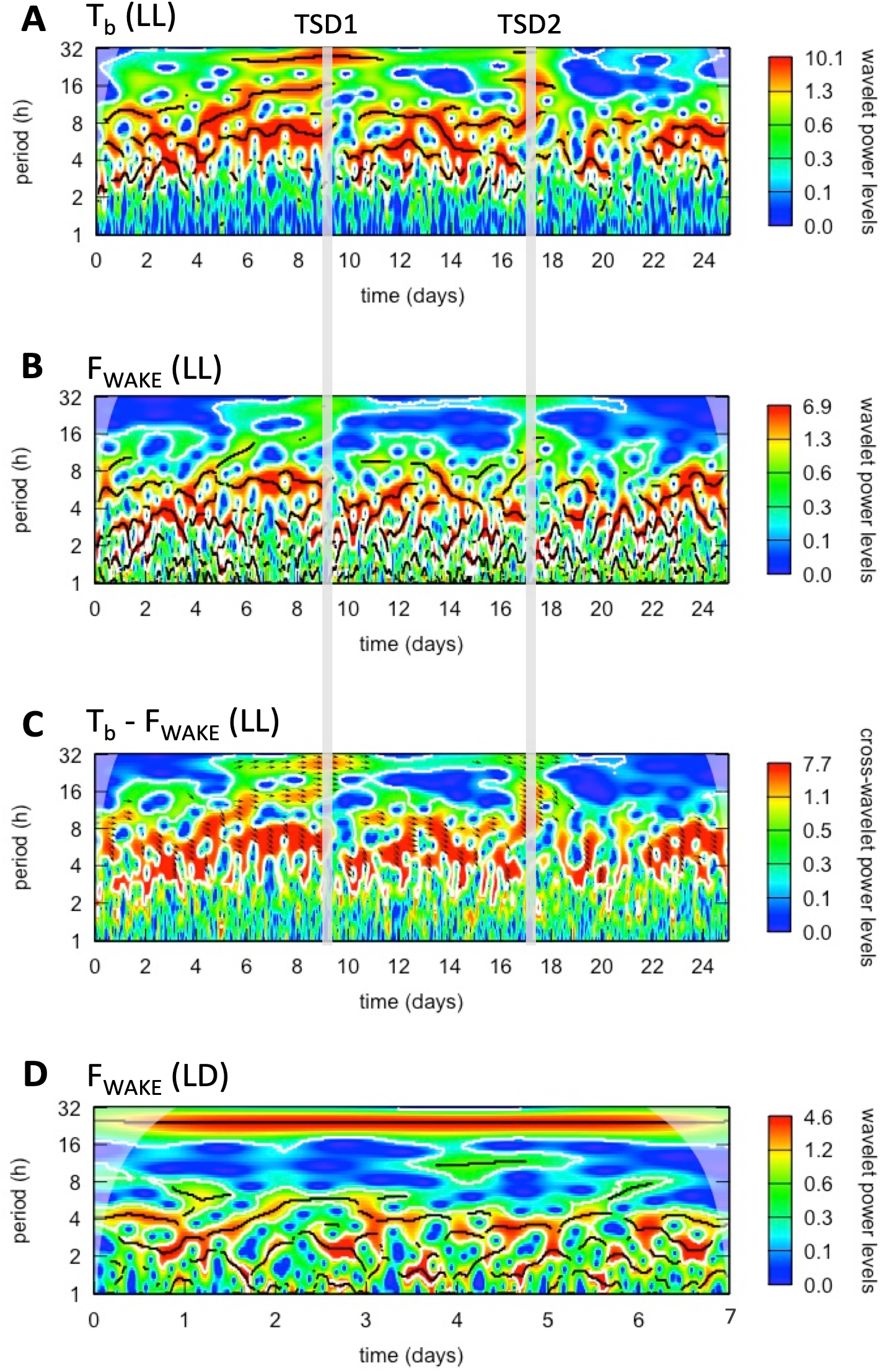
Wavelet power spectra over a complete study in a representative rat (SD4 group) under prolonged constant light (LL, panels **A-C)**, and a different rat entrained to a 12:12 h light-dark cycle (LD, panel **D**). In all panels, white contour lines denote p-values (calculated using 100 simulations) of 0.1. Power ridges (peaks across time) are indicated (in panels **A**,**B** and **D** only) by black lines on a red background of high power. Shaded regions in the top corners indicate unreliable zones due to border effects. **A**. Wavelet power levels in the body temperature (Tb) time series (mean Tb, °C, per 10 min time bin) in LL. **B**. Wavelet power levels in the F_WAKE_ time series (fractional content of WAKE per 10 min time bin) in LL. **C**. Cross-wavelet power showing periods of joint power in the bi-variate T_b_-F_WAKE_ time series in LL. The arrows indicate phase angles between the two variables. Horizonal right arrows indicate that the variables vary in phase at the periods highlighted, and the slight downward orientation indicates a small phase lag (T_b_ follows F_WAKE_ by approximately 2 time bins). **D**. Wavelet power levels in the F_WAKE_ time series (fractional content of WAKE per 10 min time bin) in LD.

Fig 1D shows an example wavelet spectrum of the F_WAKE_ time series over 7 BL days in a representative animal in LD. As expected, the diurnal 24 h rhythm is a prominent peak across the entire record. Also evident are ultradian periodicities in the 2 – 6 h range, and like the LL rats, this ultradian oscillation was non-stationary. There was a tendency for wavelet power to decrease during the early L phase of each day in LD entrained rats.

Fig 2 shows the mean (± 95% CI) phase durations of BL ultradian rhythms across circadian time in LL, compared with diurnal rhythms of rats entrained to a 12:12 h LD cycle. Phase by phase analysis was conducted across 7 LD cycles and 7 full circadian cycles in LL.

**Fig. 2.**
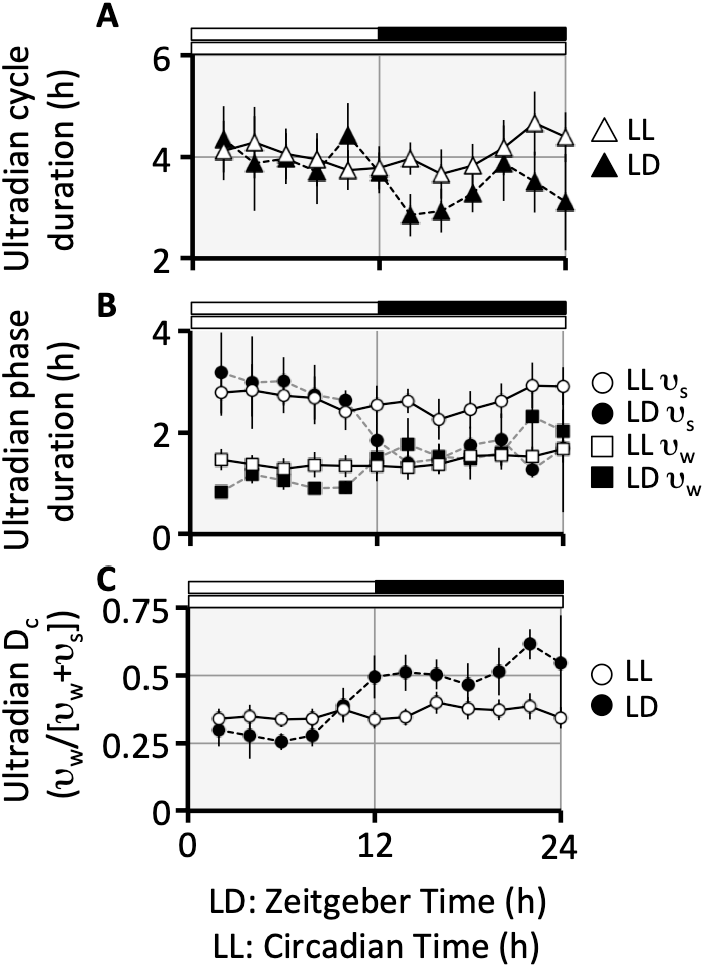
Ultradian cycle duration (A), ultradian phase durations (B) and ultradian duty cycle (C) as a function of time of day. Data are plotted as mean (± 95% CI) for n = 15 rats in LL (white symbols and solid lines) and n = 10 rats in 12:12 h LD (black symbols and dotted lines). Horizontal bars above each panel indicate light (white) and dark (black). Time scale on the abscissa is in Zeitgeber time (ZT, solar h) in LD, and in circadian time (CT, circadian h = circadian period / 24) in LL. Data were analyzed in consecutive 2 h time bins during 7 days of baseline, and are plotted at the end of each time bin. Time on the ordinate in **A** and **B** is in mean solar h. Ultradian duty cycle (conventionally defined as pulse / period in a binary cycle) is the fraction of the full ultradian cycle consisting of the υ_w_ phase.

Overall rates of state expression during BL (i.e. measured across total recording time in 7 BL days without reference to circadian or ultradian phase) were statistically different between LL (n = 15) and LD (n = 10), using Sidak’s multiple comparisons test. F_WAKE_ was 43.1 ± 1.8 % in LL and 46.8 ± 2.5 % in LD (P = 0.0167), F_NREM_ was 48.8 ± 1.8 % in LL and 41.3 ± 3.3 % in LD (P < 0.0001), and F_REM_ was 8.0 ± 0.6 % in LL and 11.9 ± 1.7 % in LD (P = 0.0096).

Thus in undisturbed BL conditions, relative to LD rats, the LL rats spent less time awake, more time asleep and, within sleep, more time in NREM sleep and less time in REM sleep.

BL data were also co-segregated by circadian (L versus D) and ultradian (υ_s_ versus υ_w_) phases. Pre-planned statistical comparisons were made using Sidak’s multiple comparisons test and the data and test results are summarized in Table 1.

**TABLE 1.** Mean (95%CI) baseline ultradian variables in the light (L) and dark (D) circadian phases. Comparison of rats kept in constant light (LL) with rats entrained to a 12:12 h LD cycle.

| Variable | LL (n=15) |  | LD (n=10) |  |
| --- | --- | --- | --- | --- |
|  | L | D | L | D |
| ultradian cycle (h) | 3.96<br>(-0.26) | 4.44<br>(-0.43) | 3.97<br>(-0.24) | 3.3*<br>(-0.18) |
| $v_s$ duration (h) | 2.61<br>(-0.21) | 2.95<br>(-0.4) | 2.84<br>(-0.2) | 1.56****<br>(-0.14) |
| $v_w$ duration (h) | 1.34<br>(-0.11) | 1.49<br>(-0.1) | 1.13<br>(-0.12) | 1.74**<br>(-0.19) |
| $D_c$ | 0.34<br>(-0.02) | 0.34<br>(-0.03) | 0.29<br>(-0.02) | 0.53****<br>(-0.04) |
| cycle count/day | 2.88<br>(-0.16) | 3.03<br>(-0.2) | 2.88<br>(-0.18) | 3.66***<br>(-0.19) |
| cycle $F_{WAKE}$ | 0.413††††<br>(-0.021) | 0.43††††<br>(-0.031) | 0.27<br>(-0.028) | 0.66****<br>(-0.036) |
| cycle $F_{NREM}$ | 0.504†††<br>(-0.019) | 0.486††††<br>(-0.024) | 0.558<br>(-0.036) | 0.272****<br>(-0.037) |
| cycle $F_{REM}$ | 0.083††††<br>(-0.007) | 0.079<br>(-0.006) | 0.172<br>(-0.022) | 0.068****<br>(-0.02) |
| $v_s F_{WAKE}$ | 0.201††††<br>(-0.022) | 0.211††††<br>(-0.023) | 0.147<br>(-0.023) | 0.396****<br>(-0.03) |
| $v_s F_{NREM}$ | 0.682†<br>(-0.02) | 0.671††††<br>(-0.017) | 0.646<br>(-0.035) | 0.475****<br>(-0.042) |
| $v_s F_{REM}$ | 0.117††††<br>(-0.01) | 0.112<br>(-0.009) | 0.207<br>(-0.027) | 0.13***<br>(-0.036) |
| $v_w F_{WAKE}$ | 0.824††††<br>(-0.02) | 0.855*†††<br>(-0.023) | 0.578<br>(-0.034) | 0.897****<br>(-0.029) |
| $v_w F_{NREM}$ | 0.158††††<br>(-0.018) | 0.13*†††<br>(-0.022) | 0.335<br>(-0.03) | 0.09****<br>(-0.028) |
| $v_w F_{REM}$ | 0.018††††<br>(-0.003) | 0.016<br>(-0.004) | 0.086<br>(-0.017) | 0.013****<br>(-0.005) |

Sidak's multiple comparisons test was used to control familywise error rate.
Within-group L v D: \*, $P < 0.05$ , \*\*, $P < 0.01$ , \*\*\*, $P < 0.001$ , \*\*\*\*, $P < 0.0001$ .
Between-group LL v LD: †, $P < 0.05$ , †††, $P < 0.001$ , ††††, $P < 0.0001$ .
LD data are a reanalysis of previously published results [37]

In Figs 2 and 3, data were grouped into 2 h time bins across the circadian cycle according to the time of onset of each ultradian cycle or phase. Systematic diurnal patterns were evident in LD that were absent in LL (Table 1). Ultradian cycle (Fig 2A) and phase (Fig 2B) durations, and ultradian duty cycle (Fig 2C) were significantly different between L and D in LD entrained rats, but not in LL animals. These ultradian timing variables changed little as a function of circadian time in LL (Fig 2), but when averaged across the entire circadian phase, were not statistically significantly different from LD values (Table 1).

**Fig. 3.**
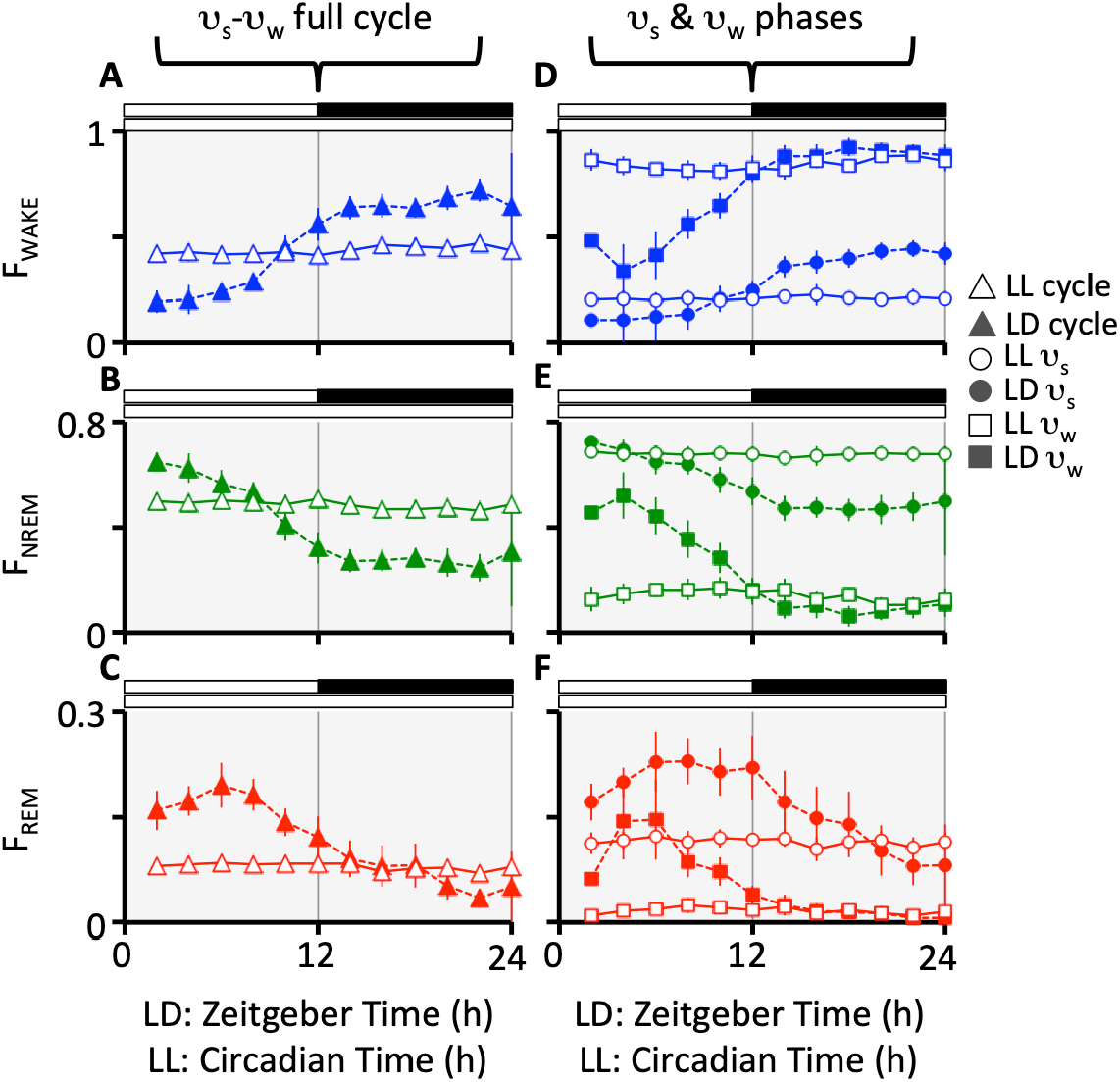
Vigilance state content as a function of time of day. **A**. Fraction of time in WAKE (F_WAKE_). **B**. Fraction of time in NREM (F_NREM_). **C**. Fraction of time in REM (F_REM_). Panels **A-C** show values calculated on a full ultradian cycle basis. Panels **D-F** show values for each of the υ_s_ and υ_w_ ultradian phases separately. Data (for 7 baseline days) were calculated for each ultradian phase and cycle and then values alotted to a given 2 h window according to time of onset of the phase/cycle. See Fig 2 caption for description of timescale. Solid symbols denote LD (n = 10) and white symbols denote LL (n = 15).

Fig 3 illustrates the circadian patterns of state expression on a full ultradian cycle-by-cycle basis (Fig 2A,B,C) and for each of the two ultradian phases separately (Fig 2D,E,F). In LD entrained rats there were highly significant differences between L and D diurnal phases in all three states (Table 1). Partitioning the data between ultradian phases revealed that rates of state expression varied in both υ_s_ and υ_w_ ultradian phases across the diurnal cycle in LD[20]. F_WAKE_, F_NREM_ and F_REM_ each differed significantly between L and D in both ultradian phases in LD (Fig 3D,E,F). Furthermore, whilst the overall circadian pattern was similar, the diurnal variation was quantitatively more pronounced in the υ_w_ ultradian phase than in the υ_s_ phase, especially during the L phase of the day. Thus, the υ_w_ – υ_s_ differences in state expression (which can be considered to be an index of ultradian amplitude) were variable and lower overall in the L phase than in the D phase under entrained LD cycles. This pattern was absent in LL rats.

In LL rats, there were no statistically significant differences between subjective L and subjective D for any state, both when measured as a function of complete ultradian cycles, and when measured in the υ_s_ ultradian phase only (Fig 3, Table 1). However, in the υ_w_ phase, there were small, statistically marginal, differences between L and D in both F_WAKE_ and F_NREM_, but not in F_REM_. This accounts for the residual circadian rhythm detected by wavelet analysis (Fig 1).

### 3.3. Sleep and wakefulness during TSD

There were no statistically significant differences between first and second TSD in any of the variables tested (duration of the TSD interval, content of WAKE, NREM and REM, rate of sleep-wake transitions, and mean bout durations of WAKE, NREM and REM), as determined by repeated measures ANOVA.

As intended, TSD durations differed between the three groups (F_(2,27)_ = 16.1, P < 0.0001), however, due to variability resulting from spontaneous wakefulness before and after stimuli, the Holm-Sidak multiple comparisons test indicated that whilst SD2 deprivations differed significantly from both SD4 (P = 0.0014) and SD6 (P < 0.0001), the difference between SD4 and SD6 was statistically non-significant (P = 0.099); SD2 = 196.4 ± 49.1 min, SD4 = 352.1 ± 87.4 min, SD6 = 421.8 ± 51.8 min. Proportion of time awake was SD2 = 0.894 ± 0.027, SD4 = 0.925 ± 0.042, SD6 = 0.918 ± 0.021. Proportion of time in NREM was SD2 = 0.091 ± 0.025, SD4 = 0.068 ± 0.036, SD6 = 0.069 ± 0.016. Proportion of time in REM was SD2 = 0.016 ± 0.005, SD4 = 0.007 ± 0.006, SD6 = 0.013 ± 0.0086.

### 3.4. Post-TSD rebound responses

#### 3.4.1. Preliminary analysis of rebound

Before the full formal analysis, an exploratory examination of post-TSD responses was undertaken using the approach described previously[10]. Briefly, cumulative time awake was obtained for each trial of each animal by summation of WAKE epochs over time of experiment encompassing 3 days before TSD, the day of TSD and 3 days following TSD. The resulting time series was detrended by least squares linear regression of the pre-TSD portion of the data. This regression line was extrapolated over the subsequent TSD and post-TSD sections and subtracted from the data to yield net cumulative WAKE (t_WAKE_) across each full TSD trial. We found the responses to be highly variable, both within and between animals, with no apparent monotonic rebound in 11 trials (see Fig 4). To distinguish response patterns, we used the Akaike Information Criterion[49] (AICc) to compare linear versus exponential least square regression of REC data. This revealed that in 3 animals (1 in the SD2 group and 2 in the SD4 group), there was no detectable monotonic rebound response in both of their trials, and in 5 other animals (2 in SD2, 2 in SD4 and 1 in SD6), one of the two TSD trials lacked a monotonic rebound. That is, only 19 of 30 trials exhibited the expected monotonic return of t_WAKE_ (and hence t_SLEEP_) toward baseline, whereas 11 trials were followed by linear t_WAKE_ that appeared to represent a new baseline, some increasing, some decreasing and others unchanged relative to pre-TSD BL. Since the objective of the study was to examine ultradian phase-dependent state transition characteristics during rebound, we treated these “rebound” (monotonic) and “non-rebound” (linear) trials separately in further analyses.

**Fig. 4.**
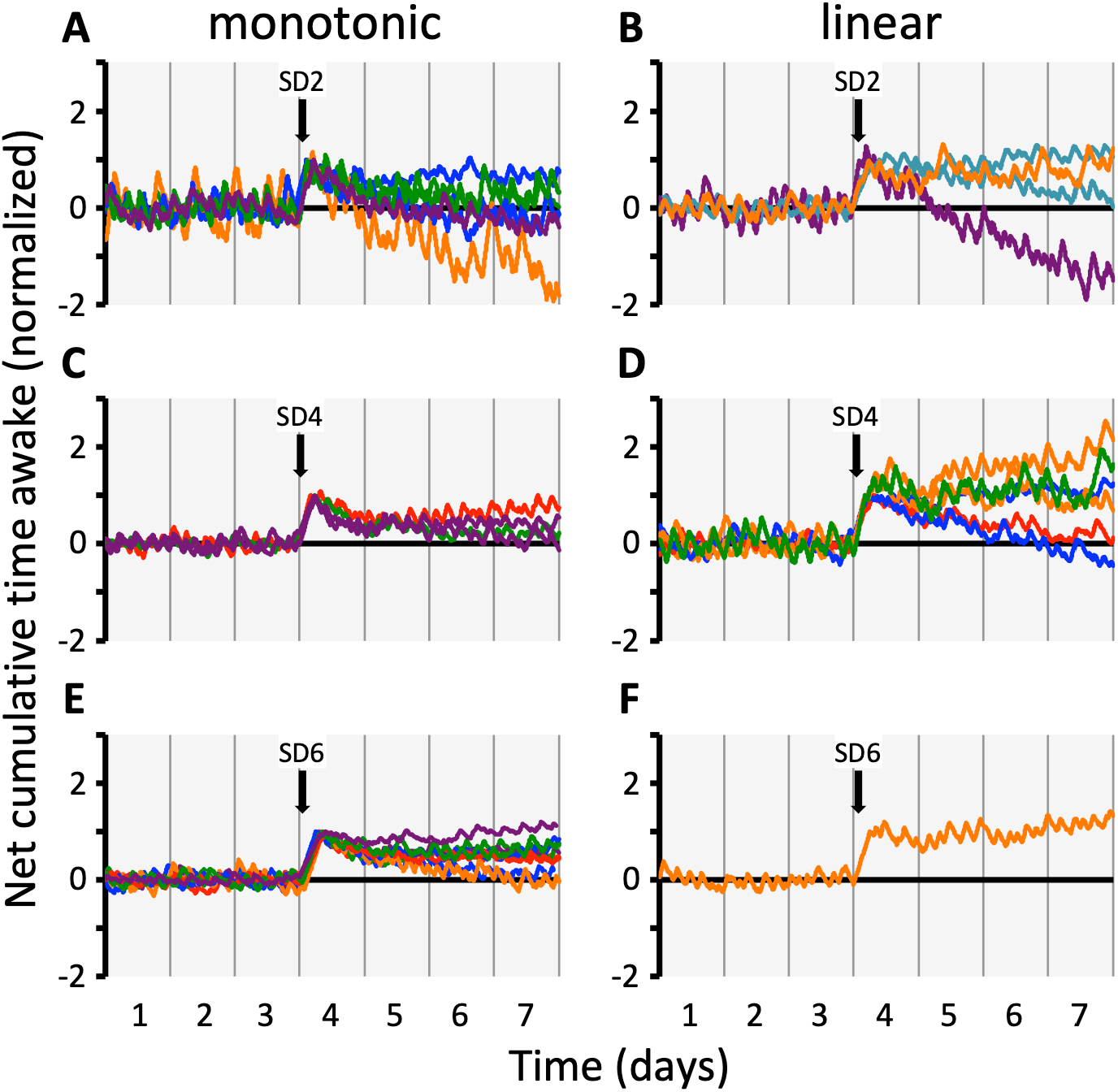
A preliminary survey of rebound responses in 3 groups (n = 5 each) of rats in LL. Each rat was exposed to two sequential 2 h (SD2; **A**,**B**), 4 h (SD4, **C**,**D**) or 6 h (SD6, **E**,**F**) episodes of total sleep deprivation (TSD) by gentle handling. Trials are aligned so that TSD onset is at the beginning of day 4 in this Figure (black arrows). Curves depict net cumulative time awake (= additive inverse of net cumulative time asleep) spanning 3 days before (pre-TSD baseline), the day containing TSD and 3 days after (post-TSD recovery). Data are normalized so that all curves are shown relative to the magnitude of the TSD-induced WAKE excess (max excess = 1 on the ordinate scale). Horizontal dashed lines indicate extrapolated pre-TSD baseline. Trials exhibiting the expected rebound profile of monotonic return to a new baseline are shown in the panels to the left for each group (**A**,**C**,**E**). The panels on the right (**B**,**D**,**F**) illustrate “atypical” linear rebound profiles. These latter trials were followed by long-term change in rates of state expression (i.e. “new” baseline slopes) that did not exhibit any identifiable correlation with baseline or TSD variables. As can be seen, in some cases net WAKE increased and in others it decreased, even within the same animal across the two trials (see traces with matching colours for examples of individual animal trials). Ultradian rhythms are clearly apparent throughout all trials, including the early stages of post-TSD rebounds – thereby refuting our previous prediction of extended υ_s_ phase during rebound[19].

#### 3.4.2. Conventional analysis of monotonic rebound

Fig 5A,B,C shows the pooled rebound as a function of time of post-TSD recovery in each group of rats (SD2, n = 6 trials; SD4, n = 4 trials; SD6, n = 9 trials). The curves show exponential regression lines and symbols denote mean ± 95% CI. Time constants (h) of the exponential rebound responses were (mean ± 95% CI): WAKE, SD2 = 12.4 ± 3.1, SD4 = 5.9 ± 1.5, SD6 = 8.2 ± 1.2; NREM, SD2 = 13.8 ± 4.5, SD4 = 5.1 ± 1.4, SD6 = 8.6 ± 1.4; REM, SD2 = 11.2 ± 2.9, SD4 = 7.9 ± 2.0, SD6 = 7.5 ± 1.6. There were no statistically significant differences in rate of rebound between groups in any state (Kruskal-Wallis test; all P > 0.8). Combining data across groups, there were no statistically significant differences in rate of rebound between states (Kruskal-Wallis test, P = 0.94). These data show that the rebounds are more than 95% complete (i.e. > 3 time constants) within 1.5 - 2 days of REC. Hence, the magnitude of the full rebound was taken in each trial as the mean of REC day 3 and used to calculate data shown in Fig 6A-C.

**Fig. 5.**
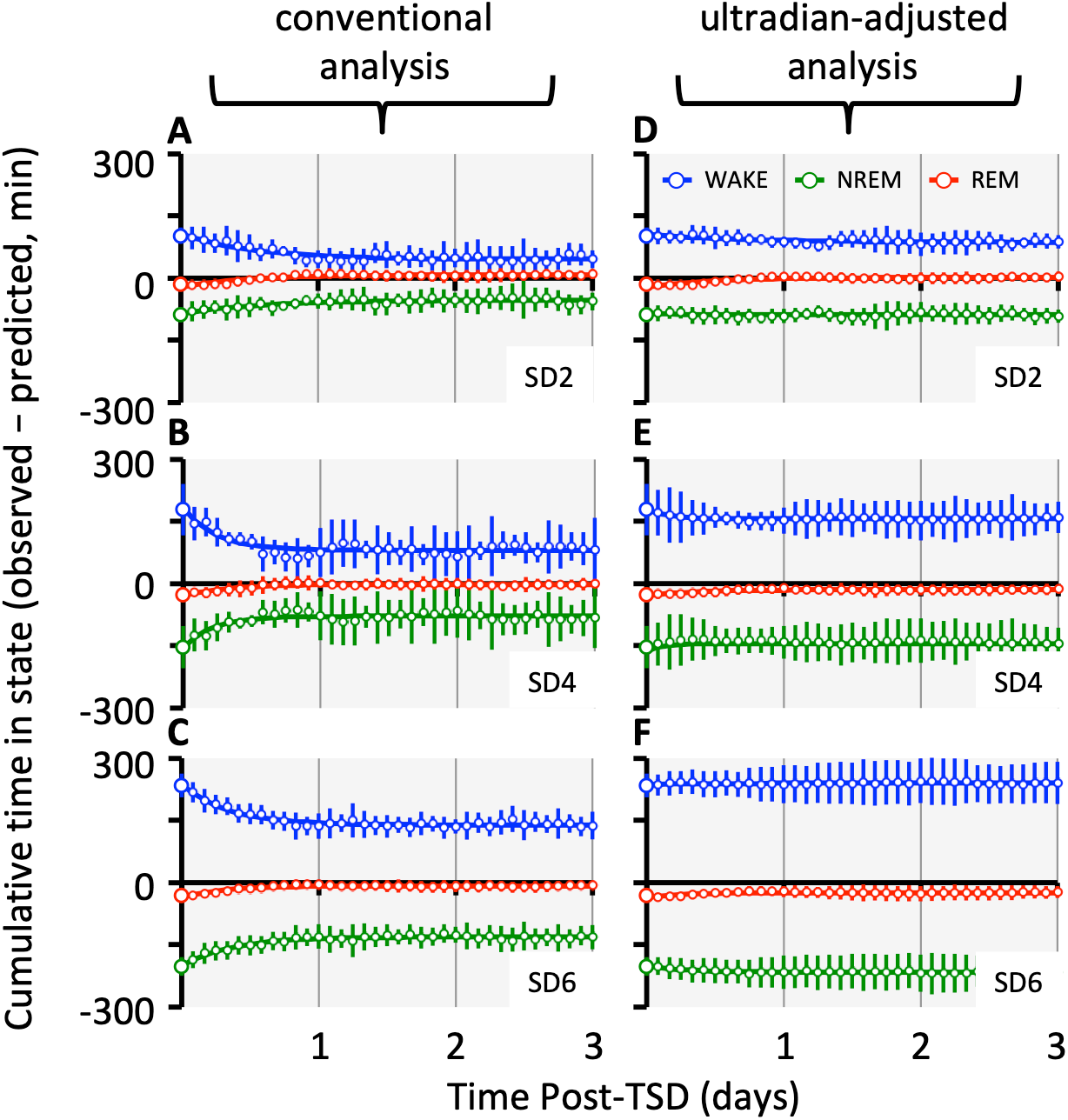
Mean (± 95% CI) monotonic rebound responses in three groups of rats in LL (SD2, n = 6 trials; SD4, n = 4 trials; SD6, n = 9 trials). In panels **A-C** data were analyzed using a “conventional” approach of calculating time in state in 36 consecutive 2 h time bins following TSD. In panels **D-F** data were reanalyzed by accounting for the contents of each of the υ_s_ and υ_w_ ultradian phases within each time bin and using phase-specific baseline mean values to calculate adjusted predictions for state content per time bin. Data are shown as the observed – predicted differences. Lines through the data are mean exponential curves obtained by least squares regression, constrained to pass through the enlarged symbols at time zero which depict TSD-induced excess/deficit values. By conventional analysis, WAKE (and therefore total SLEEP) and NREM rebounds fail to fully compensate for the TSD-induced perturbation, whereas REM rebounds do. Adjusting for ultradian phase (**D-F**) nearly eliminates the calculated rebounds of WAKE and NREM, and decreases the rebounds of REM in all groups. This indicates that ultradian timing plays an important part in determining the magnitude of the rebound.

**Fig. 6.**
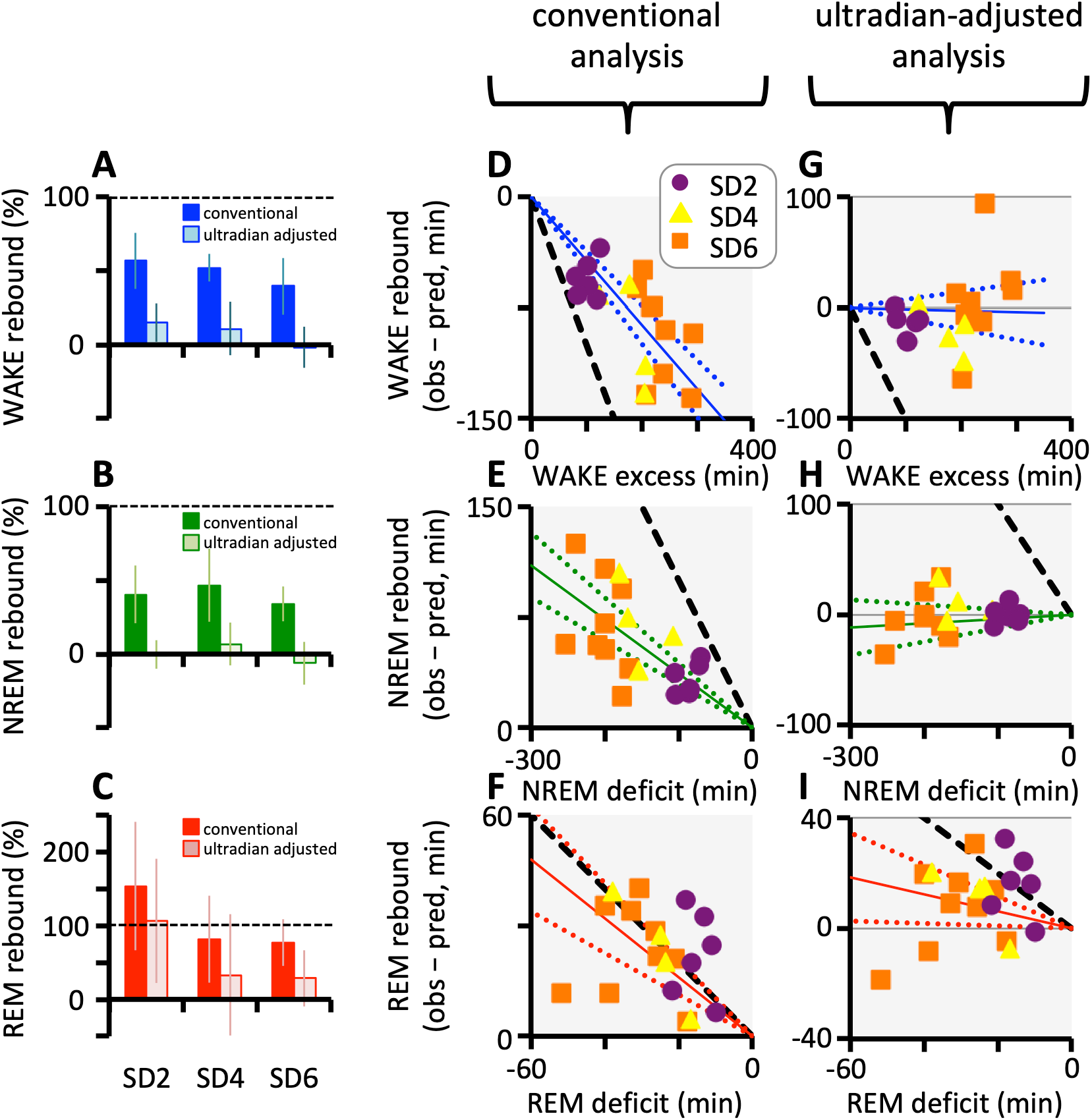
Quantification of monotonic rebounds of three groups of rats in LL (SD2, n = 6 trials; SD4, n = 4 trials; SD6, n = 9 trials). Panels A-C show mean (± 95% CI) rebounds, expressed as a percentage of the respective WAKE excess (**A**), NREM deficit (**B**) and REM deficit (**C**). Data compare rebounds calculated using the conventional approach (dark bars) and after adjustment to account for ultradian phase durations (light bars). Panels D-F depict the relationship between TSD-induced excess /deficit (abscissa) and rebound magnitude (ordinate) in three groups of rats. Dashed black lines inducate line of identity (i.e. slope = 1) representing the relation predicted for an ideal homeostat. Solid coloured lines indicate least squares linear regression through the data (groups pooled) constrained to pass through the origin. Dotted coloured lines indicate 95%CI for constrained regression. By conventional analysis WAKE (**D**), NREM (**E**) and REM (**F**) all demonstrate proportionality of response (i.e. slope significantly greater than 0) but for WAKE and NREM the proportionality constant is significantly less than 1. In contrast, the slope for REM is statistically indistinguishable from 1. After adjustment for ultradian phase, proportionality of response is eliminated (slope not different from 0) in WAKE (**G**) and NREM (**H**). Ultradian adjust resulted in greater variability of REM rebound – deficit relations (**I**) and the slope was greatly reduced but remain significantly greater than 0.

Fig 6 shows the percentage of TSD-induced perturbation that was recovered during REC in each group of rats. Data in the dark coloured bars in each of panels A,B and C illustrate the estimates obtained by using a conventional method of analysis. In all groups, WAKE (Fig 6A) and NREM (Fig 6B) rebounds failed to compensate fully for the respective TSD-induced perturbations of cumulative times in state, with relative rebounds ranging from approximately 40 – 56% among groups. In contrast, REM sleep (Fig 6C) exhibited a more complete rebound, with group mean relative rebounds ranging from 77% to 150%.

In a homeostatic control system, the controlled variable is predicted to vary (within limits) in direct proportion to the error induced by TSD. Thus, it is expected that, at least for small perturbations of the system, there will be a linear relation between the magnitudes of state rebound and TSD-induced excess or deficit, with the regression line passing through the origin (i.e. y = mx). In an ideal system, the slope of the relation (m) will be 1. Fig 6D,E,F demonstrates that, using a conventional method of analysis, the slopes of the origin-constrained linear regression between excess/deficit and rebound were significantly different from zero for WAKE, NREM and REM (all P < 0.0001). However, there was substantial scatter in the data, and in the cases of WAKE and NREM (Fig 6D,E), the slopes were also highly significantly different (both P < 0.0001) from 1. In contrast, REM rebounds, while also highly variable, did more closely approximate full compensation (Fig 6F), with a slope that was (marginally) not statistically significantly different from unity (P = 0.0915).

#### 3.4.3. Ultradian phase-adjusted analysis of rebound

In the conventional analysis described above, data were quantified in non-overlapping 2 h time bins. Since the ultradian period is highly variable on a cycle-by-cycle basis[6], it would be expected that there was a highly variable content of each ultradian phase in any given time bin across the REC interval. For this reason, we recorded the fractional content of υ_w_ and υ_s_ within each 2h bin, and found a systematic decrease (below BL) in the fractional content of υ_w_ in time bins at the beginning of REC (Fig 7A,B,C). The magnitude of this transient response was greatest in SD6 and smallest in SD2 animals, and within groups the response followed a monotonic return to BL levels over a time course matching the monotonic rebound in cumulative state (compare Fig 5A,B,C with Fig 7A,B,C). To investigate the potential impact of this ultradian timing response on the calculated state rebounds, we repeated the analysis after adjusting predicted REC values to account for ultradian phase content in each time bin.

**Fig. 7.**
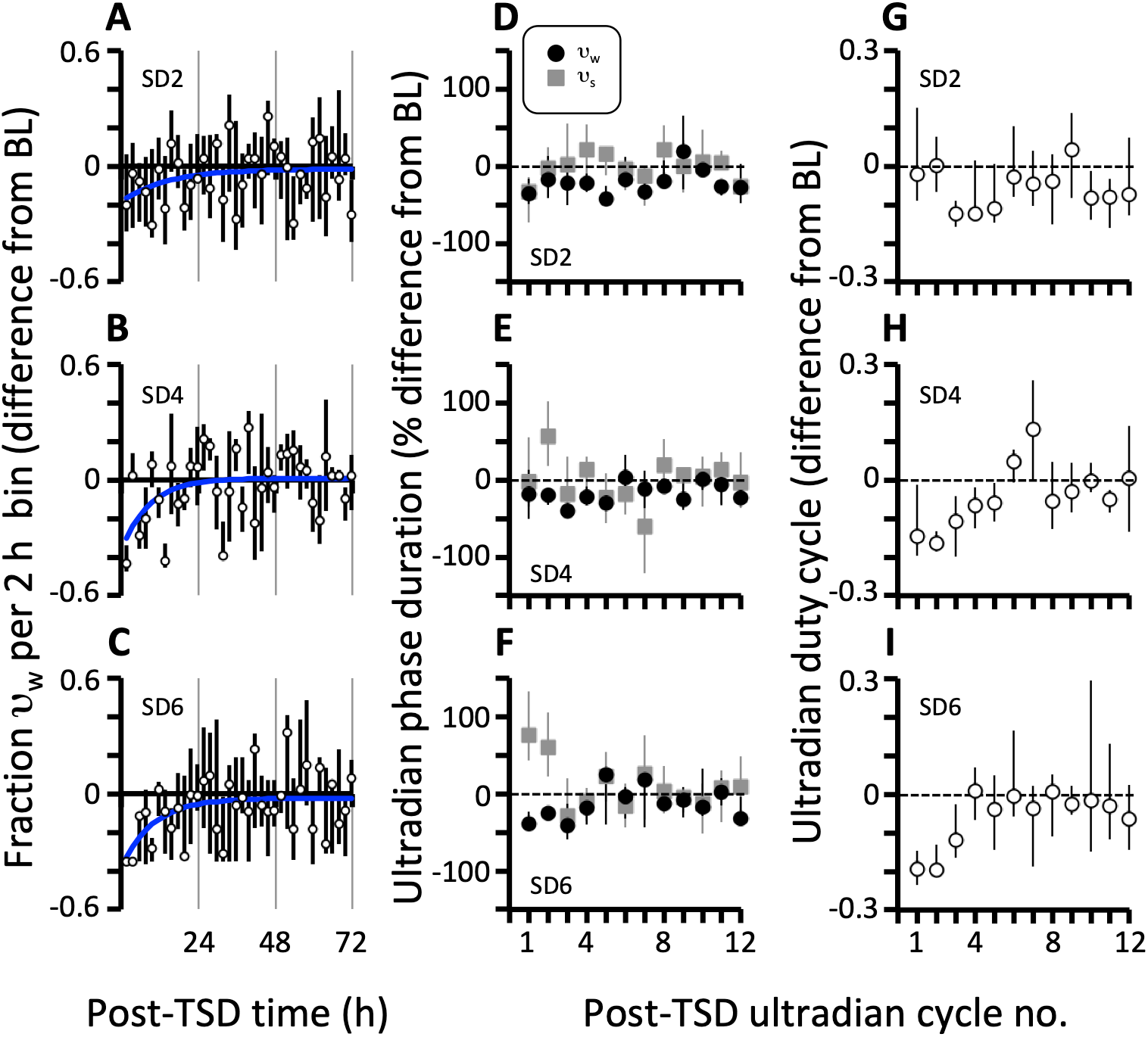
Three measures of ultradian phase responses following TSD in three group of rats (SD2, panels **A**,**D**,**G**; SD4, panels **B**,**E**,**H**; SD6, panels **C**,**F**,**I**). Data are presented as median (± inter-quartile range). In panels **A-C**, fractional content of υ_w_ phase per 2 h time bin, expressed as a difference from mean BL value. Exponential trendlines (blue) are included to illustrate the systematic suppression of υ_w_ after TSD and its monotonic return to BL values over the course of the first 1-2 days of REC. In panels **D-F**, ultradian phase durations are presented as percent change from BL values, for the first 12 ultradian cycles during post-TSD recovery. In panels **G-I**, corresponding ultradian duty cycle (υ_w_ duration / cycle duration) is given as difference from BL values. The data shown in the first and third columns represent two ways of displaying the same phenomenon, and both are a result of the slightly divergent responses of the two ultradian phases shown in the middle column.

Figs 5 and 6 demonstrate that in all three groups the rebound responses of WAKE and NREM were almost entirely abolished by taking changes in ultradian phase durations into account. In contrast, a small rebound in REM sleep remained. Fig 6G,H shows an absence of proportionality between rebounds and corresponding WAKE excess/NREM deficit (i.e. slopes are not different from zero). In contrast, following adjustment for ultradian timing there was a reduced, but nevertheless statistically significant (one-sample t test: P = 0.0228) REM rebound in the SD2 group (Fig 6C). However the ultradian-adjusted REM rebound was not statistically significant in groups SD4 and SD6 (P = 0.298 and P = 0.1221, respectively). Fig 6I shows that, after accounting for ultradian phase the magnitude of REM rebound was linearly related to REM deficit (i.e. slope ≠ 0, P = 0.0261), albeit at a much reduced slope and with substantial scatter in the data. The above results suggest that ultradian timing plays an important role in mediating the post-TSD rebounds of all three states. Specifically, a post-TSD change in the proportion of time in υ_s_ and υ_w_ per 2 h time bin gave rise to almost all of the rebounds of WAKE and NREM, and a substantial part of the REM rebound. The part of the REM rebound that was not accounted for by this timing mechanism was likely attributable to changes in state transition probabilities within one or both of the ultradian phases. To gain further insight we re-examined the responses on an ultradian phase-by-phase basis in the “rebounder” subgroup of trials.

#### 3.4.4. Ultradian phase-specific analysis of rebound

Fig 7D,E,F illustrates deviations from mean BL values with respect to durations of the first 12 post-TSD ultradian υ_s_ and υ_w_ phases in the three groups of rats. In all three groups, there was a decrease in υ_w_ durations below BL over the first 4 ultradian cycles, and less consistent responses in υ_s_. An important consequence of this differential pattern of responses between υ_w_ and υ_s_ was a transient net decrease in the fraction of each ultradian cycle consisting of the υ_w_ phase (i.e. the “duty cycle”, D_c_), most notable in the SD4 and SD6 groups (Fig 7G,H,I).

Fig 8 shows the mean (± 95% CI) normalized (%) rebound response in sequential post-TSD ultradian cycles for WAKE, NREM and REM. Note that incomplete rebounds in WAKE (Fig 8A,B,C) and NREM (Fig 8D,E,F) are evident when calculated for full ultradian cycles, but not in either of the υ_w_ or υ_s_ phases individually. In contrast, a complete REM rebound was confirmed when calculated on a full ultradian cycle basis, and a partial REM rebound was present in the υ_s_ ultradian phase alone (Fig 8G,H,I). There was no detectable REM rebound in the υ_w_ phase.

**Fig. 8.**
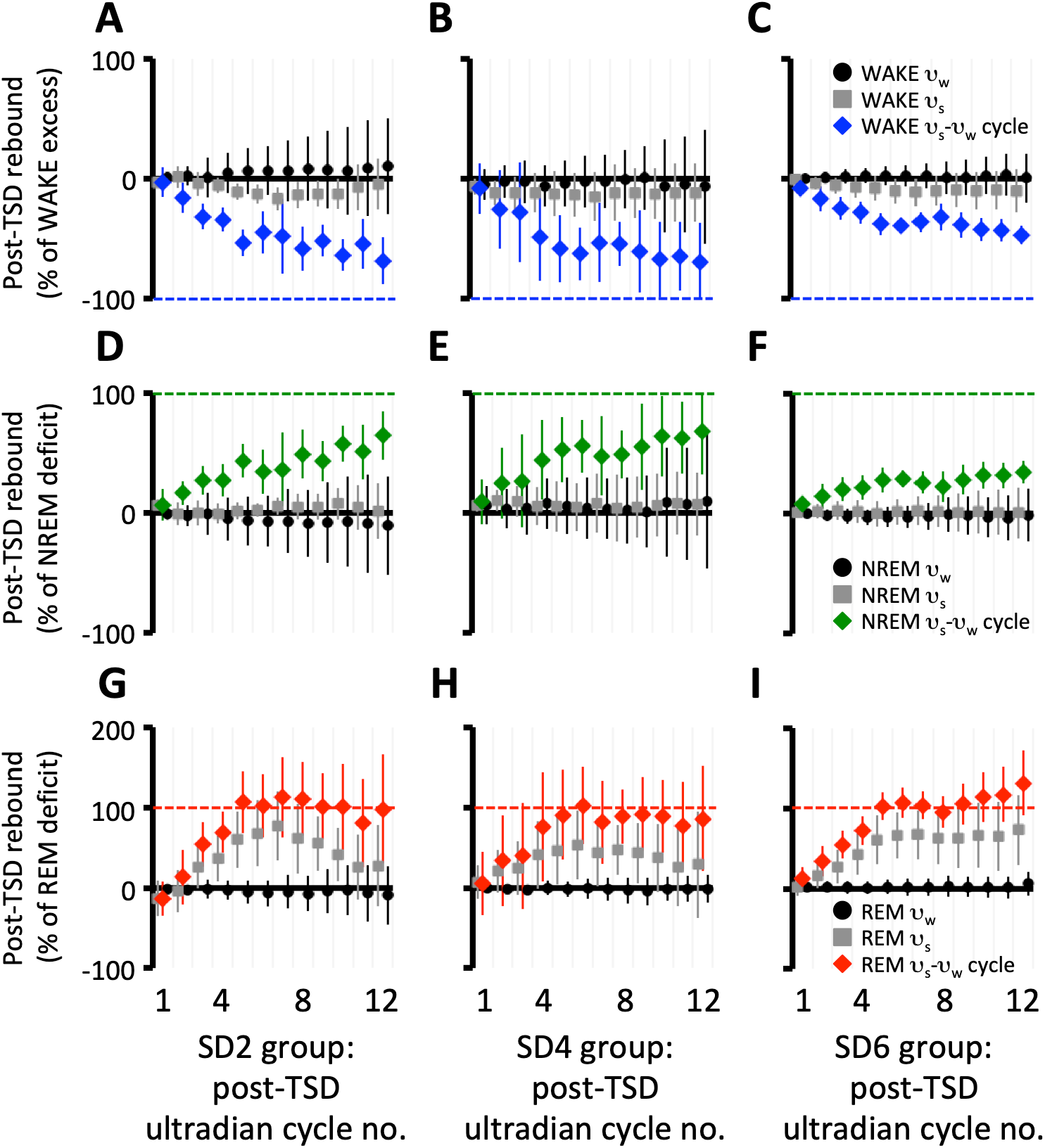
Post-TSD rebounds in 3 groups of rats: SD2, 2 h deprivation, n = 6 trials (panels **A**,**D**,**G**); SD4, 4 h deprivation, n = 4 trials (panels **B**,**E**,**H**); SD6, 6 h deprivation, n = 9 trials (panels **C**,**F**,**I**). Rebounds represent cumulative time in state and were calculated on an ultradian cycle-specific basis in each animal, and normalized as percent difference from baseline (BL) prediction. Rebound was assessed over the first 12 ultradian cycles following TSD. The scale on the abscissa represents ultradian cycle number and is not linearly proportional to time (see Fig 7). **A-C**. Ultradian cycle-by-cycle rebounds for WAKE. Calculations on a full cycle basis (blue) yield a significant but incomplete rebound in WAKE. Since change in WAKE = -change in total sleep, these WAKE rebounds are quantitatively identical (but opposite sign) to corresponding rebounds of total sleep. Note the absence of WAKE rebound in either of the υ_s_ (grey) and υ_w_ (black) phases alone. **D-F**. Ultradian cycle-by-cycle rebounds for NREM. Incomplete rebounds are present on a full cycle basis (green) but absent in either of the υ_s_ (grey) and υ_w_ (black) phases alone. **G-I**. Ultradian cycle-by-cycle rebounds for REM. Fully complete rebounds are present on a full cycle basis (red) and partially complete rebounds are present in the υ_s_ (grey) phase alone, but absent in the υ_w_ (black) phase alone. This indicates that important intra-phase adjustments contribute to the REM rebound during the υ_s_ phase, but not the υ_w_ phase of the ultradian cycle.

The absence of any rebound in the υ_w_ phase implies a lack of response at the state transition level within that phase. In support of this, post-TSD values for bout durations, bout frequencies and probabilities of state transition trajectory were all statistically similar to corresponding BL values in υ_w_.

##### 3.4.4.1 Bout durations in the ultradian υ_s_ phase

Bout durations were little changed from BL when examined on a υ_s_ phase-specific basis (Fig 9). The only response of note was a trend toward increased NREM bout durations in the SD4 and SD6 groups early in REC (Fig 9E,F). In light of the presence of a relatively strong REM rebound in υ_s_, it was noteworthy that REM bout durations were not different from baseline in this ultradian phase (Fig 9G,H,I).

**Fig. 9.**
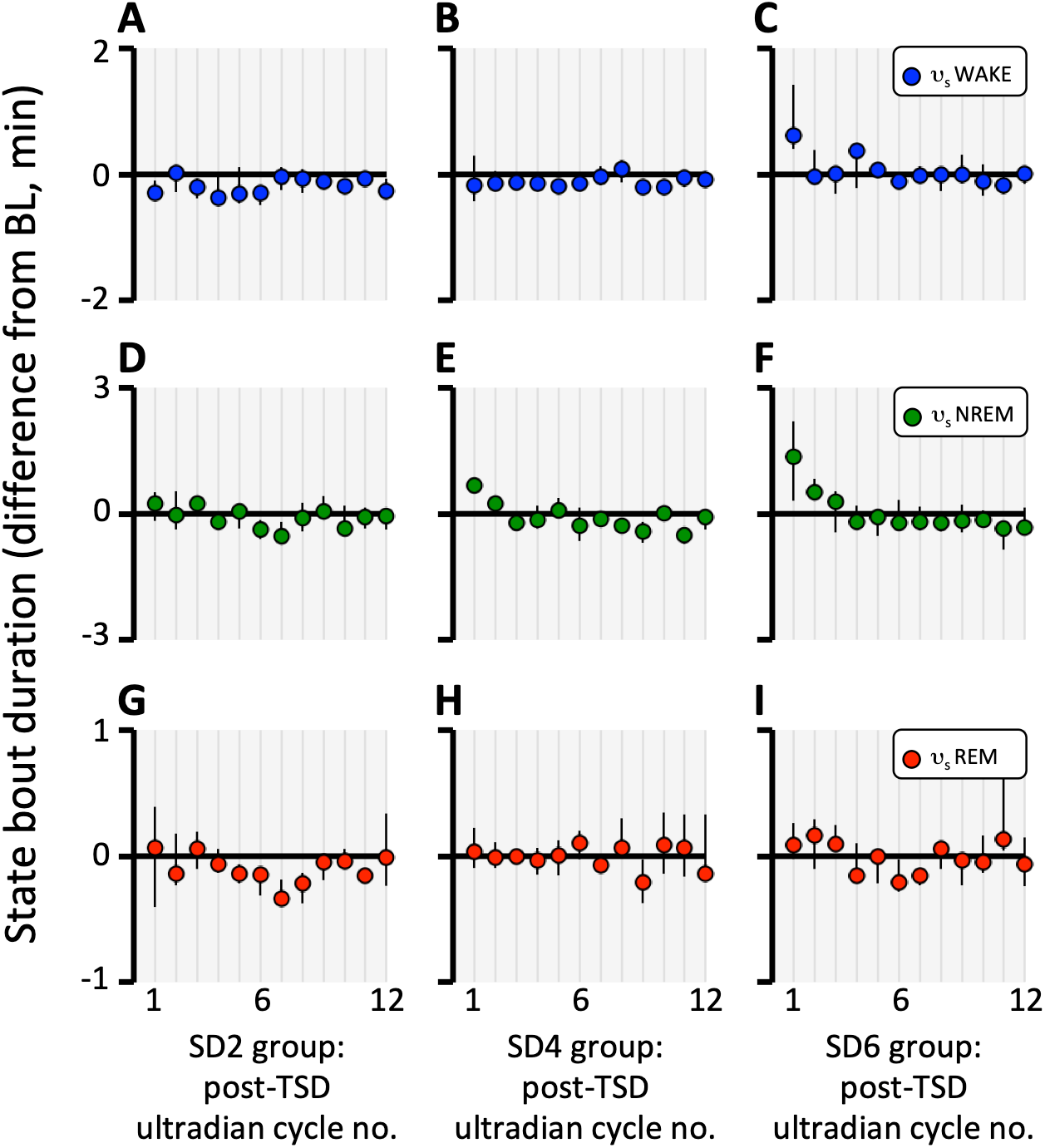
Median (± inter-quartile range) state bout durations, expressed as difference from BL, in the υ_s_ phases of the first 12 ultradian cycles following TSD. Data are shown for monotonic response trials in 3 groups of rats: SD2 (panels **A**,**D**,**G**); SD4 (panels **B**,**E**,**H**); SD6, (panels **C**,**F**,**I**). Within-phase changes in bout duration were entirely absent in the υ_w_ phase and are not shown. Responses were highly variable within and between trials and the only consistent response in the υ_s_ phase was a small transient increase in NREM bout duration in the SD4 and SD6 groups.

##### 3.4.4.2. Bout frequencies in the ultradian υ_s_ phase

As would be predicted by zero-sum considerations[37], the increase in NREM bout durations gave rise to corresponding reductions in WAKE and sleep bout frequencies (Fig 10). However, the reduction in sleep episode frequencies was due entirely to reduced NREM bout frequency, with no change from BL in REM bout frequency. This implies that the increase in F_REM_ that constitutes the partial REM rebound in υ_s_ is due to an increase in the *relative* frequency of REM bouts arising indirectly from a constant *absolute* REM frequency and decreased frequency of the other two states. This further implies that there was an adjustment in post-NREM transition probabilities that sustained a constant absolute REM frequency.

**Fig. 10.**
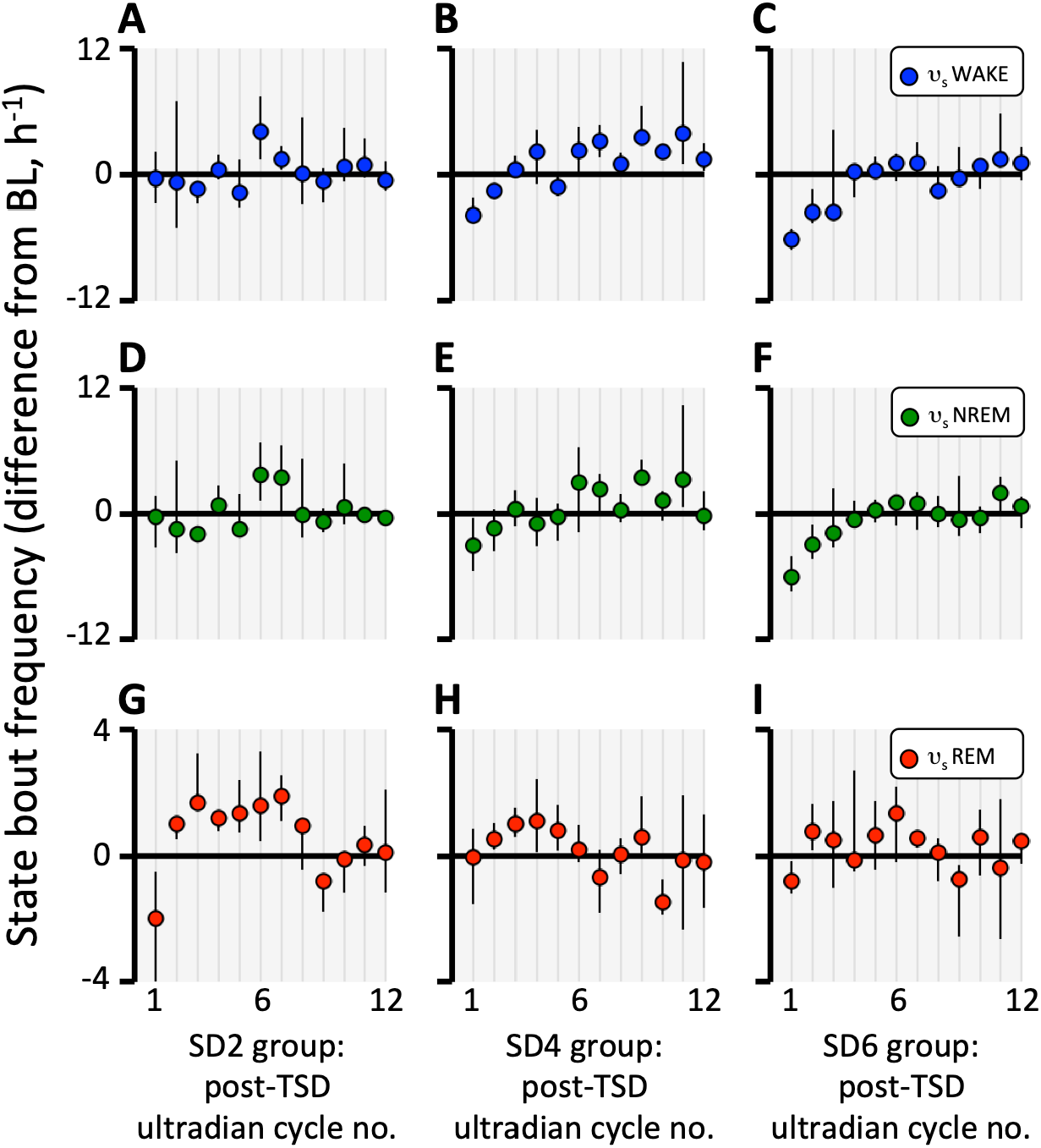
Median (± inter-quartile range) state bout frequencies (bouts / h), expressed as difference from BL, in the υ_s_ phases of the first 12 ultradian cycles following TSD. Data are shown for monotonic response trials in 3 groups of rats: SD2 (panels **A**,**D**,**G**); SD4 (panels **B**,**E**,**H**); SD6, (panels **C**,**F**,**I**). Within-phase changes in bout frequency were entirely absent in the υ_w_ phase and are not shown. The small increases in bout duration shown in Fig 9 were expected to give rise to corresponding decreases in bout frequency[37]. This was observed in the SD4 and SD6 groups for WAKE (**B**,**C**) and NREM (**E**,**F**). However REM bout frequency was sustained at BL levels (**H**,**I**).

##### 3.4.4.3 State transition trajectory in the ultradian υ_s_ phase

Fig 11A,B,C shows that the maintenance of BL-level REM frequency in the face of elevated NREM bout duration, did not involve an increase in the fraction of WAKE to REM transitions (sleep-onset REM, SOREM). Instead, this was brought about through a change in the fraction of NREM bouts that transitioned to WAKE and REM (i.e. increased tendency for NREM – REM sleep cycling and corresponding decreased tendency for NREM arousals). This is shown in Fig 11D,E,F where there was a trend for normalized probability of arousal in the ultradian υ_s_ phase to decrease transiently following TSD. The post-REM transition trajectory showed a similar but less consistent response (Fig 11G,H,I). Increased mean NREM bout durations imply increased probability of state maintenance and corresponding decrease in probability of NREM transition. Furthermore, maintenance of REM frequency implies that the decrease in P_*nw_ involved both a decrease in absolute probability of arousal (P_nw_) together with a smaller increase or no change in absolute probability of REM onset (P_nr_). These considerations suggest that the maintenance of REM frequency at BL levels during REC, which was responsible for the partial REM rebound during the ultradian υ_s_ phase (Fig 8G,H,I), was primarily an indirect result of a decrease in arousal, rather than an increase in REM “pressure” per se.

**Fig. 11.**
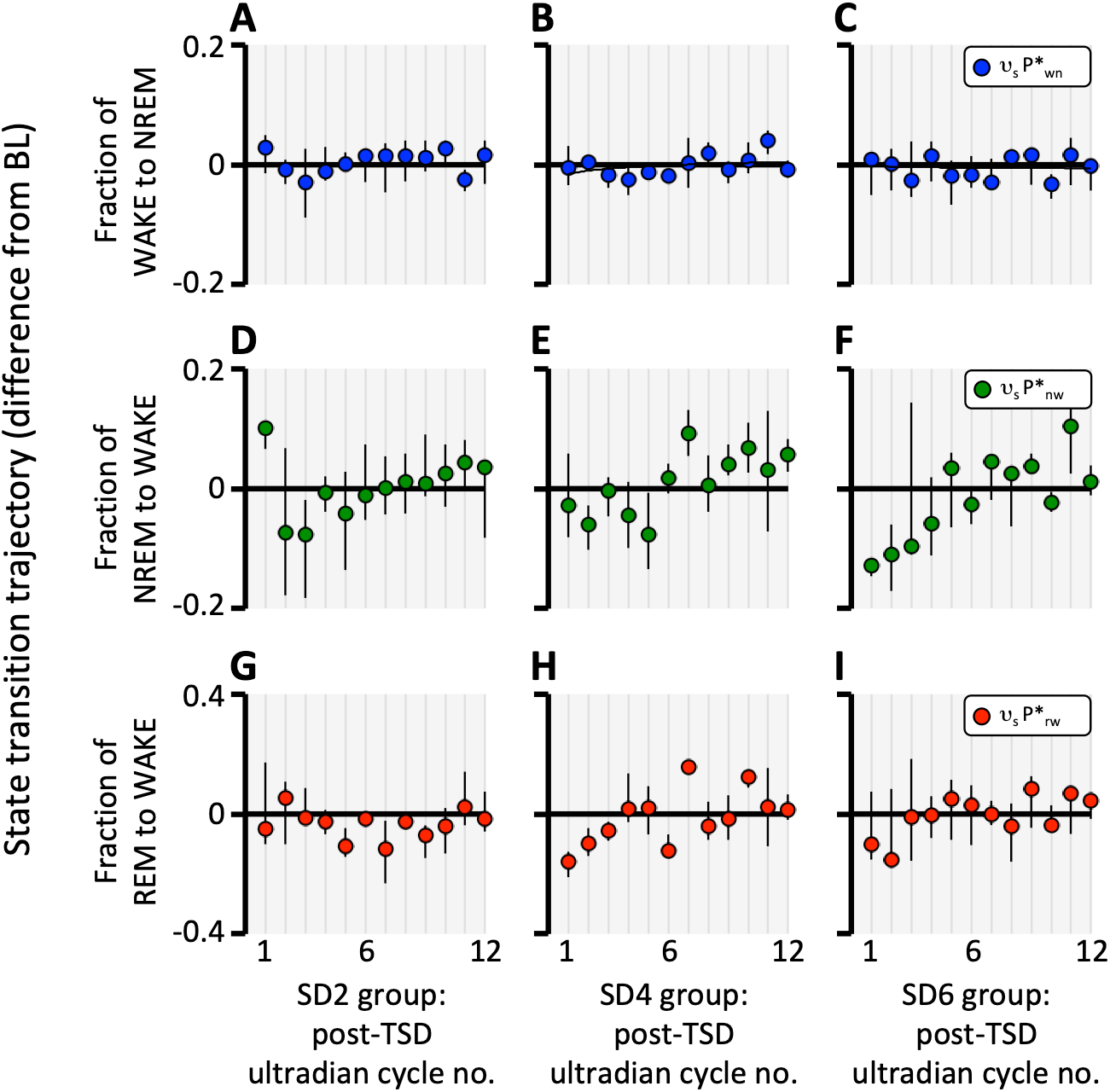
Median (± inter-quartile range) state bout transition trajectory, expressed as difference from BL, in the υ_s_ phases of the first 12 ultradian cycles following TSD. Data are shown for monotonic response trials in 3 groups of rats: SD2 (panels **A**,**D**,**G**); SD4 (panels **B**,**E**,**H**); SD6, (panels **C**,**F**,**I**). Within-phase changes in bout transition trajectory were entirely absent in the υ_w_ phase and are not shown. In a three-state system, each state can transition to one of two other states. The trajectory of these transitions can be expressed as a relative probability, estimated as the fraction of bouts in each state that transition to each of the two possible options. For WAKE, the relative probability of NREM onset 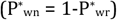 was unchanged from BL following TSD in all three groups of rats (**A-C**). For NREM, there was a marked transient decrease in the relative probability of arousal 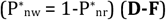 (**D-F**). Since the decrease in 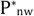 is accompanied by an increase in NREM bout durations (i.e. increase in probability of state maintenance, P_nn_) (Fig 9), then this implies that the change in absolute probability of arousal is greater than any change in absolute probability of REM onset, because P_nn_ + P_nw_ + P_nr_ = 1, and 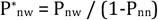. It is this change in the post-NREM transition trajectory that ensures maintained REM bout frequencies (Fig 10). Relative probability of REM arousal 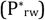 showed a similar, though less consistent trend for a transient decrease following TSD (**G-I**).

##### 3.4.4.4. Correlations in bout vs subsequent inter-bout intervals within each of the υ_w_ and υ_s_ phases

Sample sizes precluded meaningful correlation analyses on an ultradian cycle-by-cycle basis, and so we pooled data across BL and across the first 6 post-TSD ultradian cycles (because the monotonic rebound was clearly ongoing during these cycles (Fig 8)). Spearman rank correlation coefficient (ρ) was calculated, in each of BL and REC, for WAKE bout duration versus the duration of subsequent sleep episode and for REM bout duration versus the duration of subsequent inter-REM intervals (IRI).

In the case of the WAKE-sleep sequences, there were no significant correlation coefficients in either υ_w_ or υ_s_ in groups SD4 and SD6 during BL or REC. However, there was a small positive correlation in both BL and REC in the SD2 group (BL, ρ = 0.139 ± 0.064, group median P = 0.0002 and REC, ρ = 0.135 ± 0.078, P = 0.0057, respectively). Spearman’s ρ was not significantly different between BL and REC in all three groups (all P ≥ 0.158).

In the case of the REM – IRI sequence, all groups had statistically significant positive correlations in both BL and REC in the υ_s_ phase. In BL, SD2, ρ = 0.30 ± 0.06, SD4, ρ = 0.31 ± 0.10, SD6, ρ = 0.37 ± 0.07 (all P < 0.0001).There were no significant differences between BL and REC in any group (all P > 0.3). In contrast, positive ρ values were lower (< 0.2) and not statistically significant in the υ_w_ phase in both BL and REC in all groups.

##### 3.4.4.5. Ultradian phase-specific analysis of NREM EEG slow wave activity rebound

The partial rebound in NREM sleep time (t_NREM_) was evident on a per ultradian cycle basis but was not present in either the υ_w_ or υ_s_ phases (Fig 8D,E,F). According to a long-standing hypothesis[50, 51], the shortfall in cumulative t_NREM_ rebound might be offset by an increase in NREM intensity, as indicated by EEG delta power during NREM sleep (SWA_NREM_).

We confirmed that mean SWA_NREM_ was elevated above BL at the onset of post-TSD recovery in LL, and that this was proportional to the duration of TSD (Fig 12A,B,C). Interestingly, this positive rebound was present in NREM bouts in the υ_s_ phase but absent from NREM bouts in the υ_w_ phase. We further confirmed a subsequent undershoot (“negative rebound”) in SWA_NREM_, which began at approximately the fourth ultradian cycle of the rebound. This was also proportional to the duration of the TSD interval and was present in both υ_w_ and υ_s_ phases of the ultradian cycle.

**Fig. 12.**
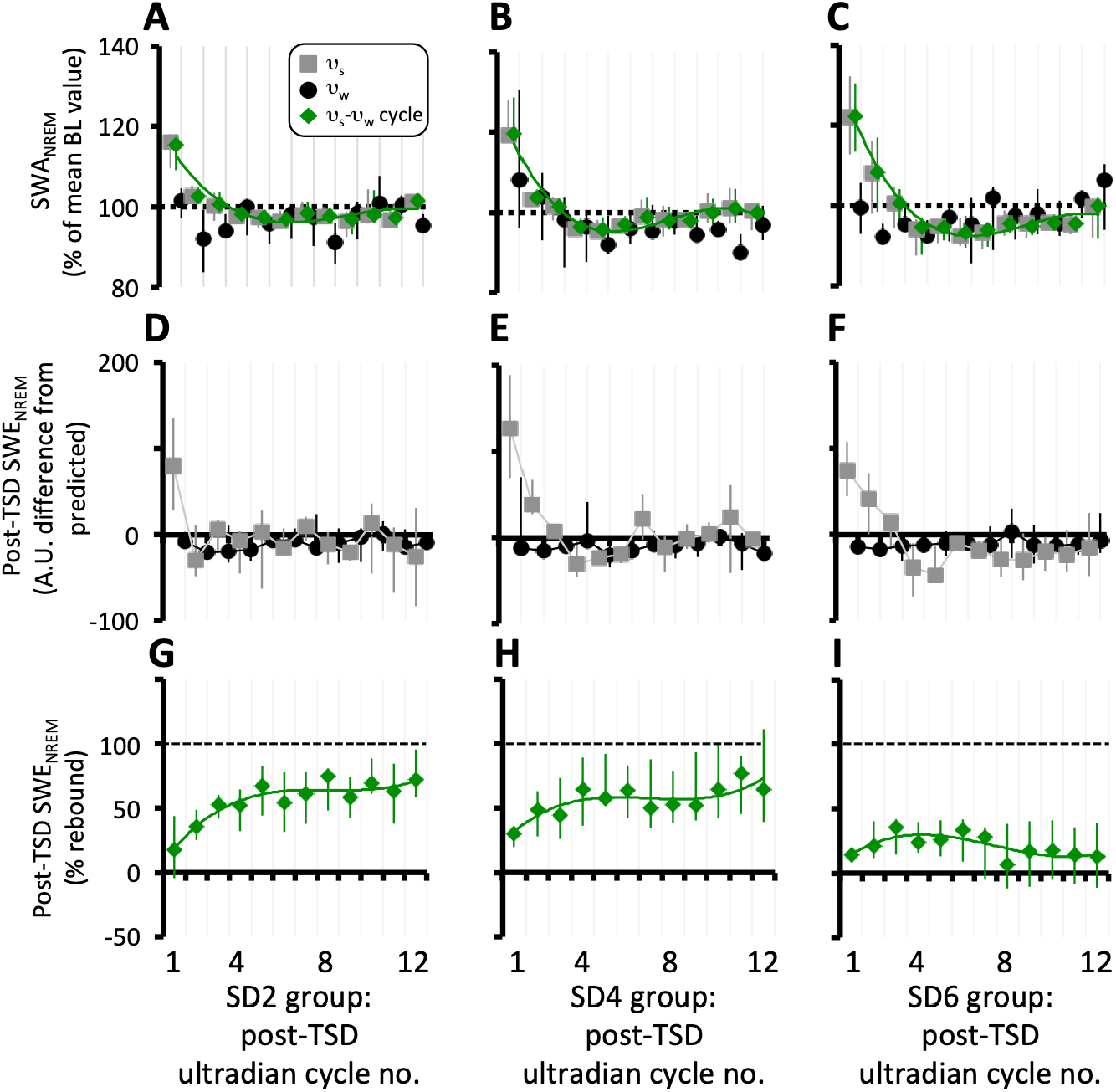
Rebound responses of putative NREM sleep “intensity”, as indexed by SWA_NREM_, and cumulative time-intensity indexed by SWE_NREM_, in υ_w_ (black), υ_s_ (grey) and full ultradian cycles (green) in 3 groups of rats SD2 (left panels), SD4 (centre panels) and SD6 (right panels). Polynomial (4^th^ order) trendlines are plotted through full cycle data for illustrative purposes only. Data depict differences from corresponding baseline values (normalized as % in first and third rows) and are plotted as median (± inter quartile range) in each of the first 12 post-TSD ultradian cycles. Panels **A-C**, show EEG slow wave activity in NREM sleep (SWA_NREM_), a putative marker of NREM sleep intensity. Full ultradian cycle and υ_s_ phase-specific responses were similar – transient positive rebound (i.e. higher than baseline) for the first 2 ultradian cycles, followed by a more prolonged but lower magnitude negative rebound (sub-baseline). In υ_w_, the positive rebound was absent but a small negative rebound was apparent. Both positive and negative rebounds appeared to be proportional to TSD duration across groups. **D-F**. Ultradian phase-specific cumulative NREM slow wave activity (slow wave energy, SWE_NREM_) calculated as the product of time in NREM state and SWA_NREM_. Data illustrate a transient positive rebound lasting approximately 3 ultradian cycles in the υ_s_ phase only. **G-I**. Post-TSD SWE_NREM_ rebounds expressed as a percentage of TSD-induced SWE_NREM_ deficit. Horizontal dashed lines indicate full rebound predicted by the delta homeostasis model [51].

The product of the two responses to TSD – time in NREM and hypothetical intensity of NREM – can be considered to be an index of the homeostatically regulated variable in NREM sleep[4]. In the present study this quantity (SWE_NREM_) was found to exhibit an approximately monotonic rebound (Fig 12D,E,F) that was not proportional to preceding TSD duration and was apparent in υ_s_ but not in υ_w_. None of the groups exhibited a full rebound in SWE_NREM_, and indeed the percent rebound in SWE_NREM_ was no greater than the percent rebound in t_NREM_ (Fig 8G,H,I).

##### 3.4.4.6. Factors influencing post-TSD baseline in linear “non-rebound” and post-monotonic “rebound” trials

As illustrated in Fig 4B,D,F, there were 11 trials in which post-TSD REC lacked the expected monotonic rebound. These trials were better fit by a linear regression than by an exponential, as determined by AICc, and were considered for the purpose of analyses to be a separate group of “non-rebounders”. It can be seen in Fig 4 that not only were these REC responses fairly linear, but that the slopes varied between trials, even within the same animal. We assume that these linear cumulative WAKE and sleep records represent slightly altered post-TSD baselines. It was noted that several of the “rebounder” trials also showed changes in baseline slope after the monotonic phase of recovery had reached completion (Fig 4A,C,E). It should be pointed out that the depiction in Fig 4 of the data as net cumulative time emphasizes such changes in slope. In fact, the actual change in rate of state expression was small in all cases, less than ±5 min/h for WAKE and NREM, and less than ±1.5 min/h for REM.

Fig 13 illustrates the observed pre-to post-TSD difference in baseline rates of state expression plotted against the difference predicted according to the corresponding changes in ultradian duty cycle. The same relation exists for both “rebounders” (white symbols) and “non-rebounders” (coloured symbols) and least squares linear regression analyses of the pooled data are illustrated. There was a linear relation, in WAKE and NREM, but no significant relation in REM. In all three states the regression lines passed through the origin (i.e. intercepts not significantly different from 0). For WAKE, the regression slope was significantly different from both 0 (slope = 0.79 ± 0.10; P < 0.0001) and 1 (P < 0.05). For NREM, the regression slope was significantly different from 0 (slope = 0.91 ± 0.14; P < 0.0001) but not different from 1. For REM the slope was not significantly different from 0 (slope = 0.10 ± 0.42; P = 0.81) and highly significantly different from 1 (P < 0.0001). Coefficient of variation (R^2^) indicated that regression lines explained 68%, 60% and 0.2% of the variance in the data in WAKE, NREM and REM, respectively. Thus, pre-to post-TSD change in ultradian duty cycle was the main factor responsible for corresponding changes in baseline rates of WAKE and NREM expression, but it had no detectable influence on any changes in the rate of REM expression.

**Fig. 13.**
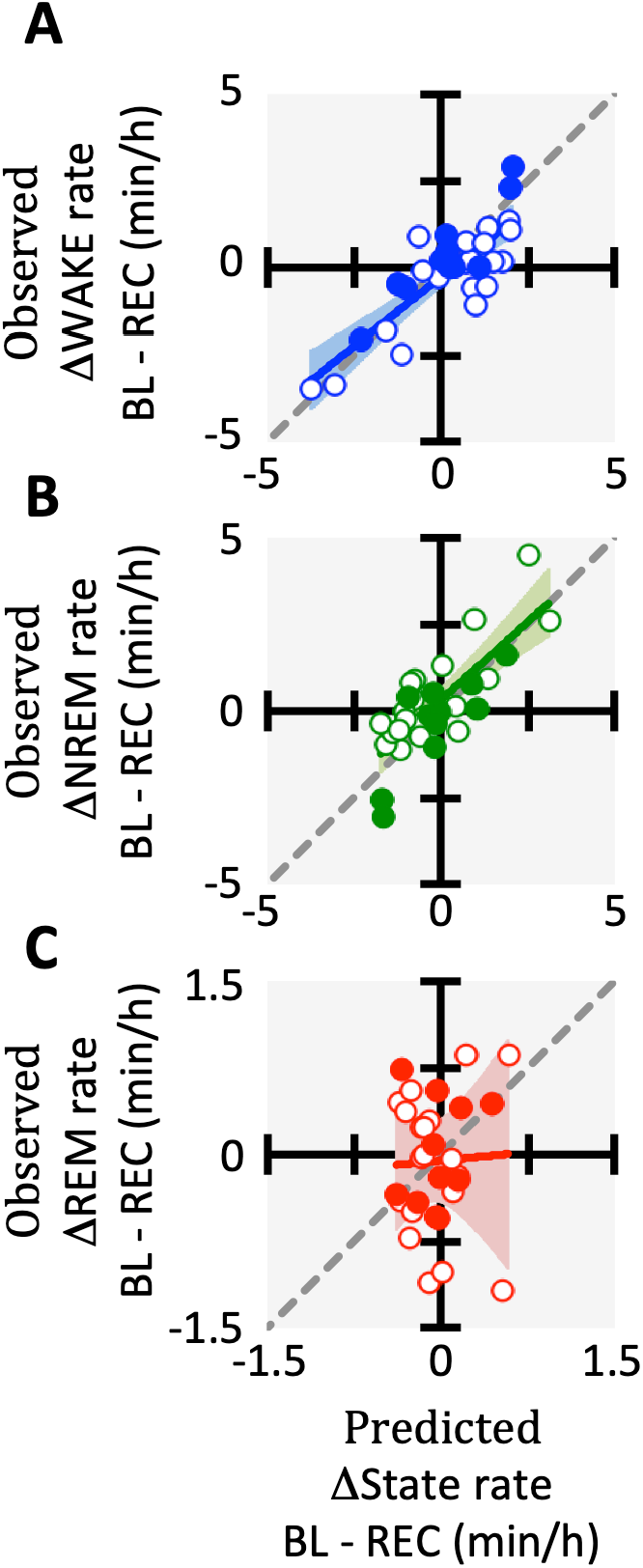
Effect of variation in ultradian duty cycle (the change in fraction of total cycle time comprising the υ_w_ phase) on baseline rate of state expression. Both axes express the change in steady baseline before vs after TSD. All trials are shown, with both “rebounders” (trials followed by monotonic rebounds, white symbols) and “non rebounders” (trials followed by linear responses, solid coloured symbols) depicted separately. Slopes were calculated on the pooled data, following completion of the monotonic response in “rebounders”. The charts deptict observed (ordinate) versus predicted (abscissa) change in rates of state expression. The prediction is based on the theoretical change resulting from the observed change in D_c_. The line of identity (observed = predicted) is shown in each case as a grey dashed line. Least squares linear regression through the data are shown as coloured lines and the 95% confidence interval of the regressions are indicated by coloured shaded regions. TSD-induced changes in baseline rates of WAKE (**A**) and NREM (**B**) are closely linked to changes in ultradian timing (i.e. D_c_). However, baseline rate of REM (**C**) expression is little changed by TSD and not affected consistently by the TSD-induced chages in D_c_.

The line of identity (slope = 1) is shown as a grey dashed line and represents the relation expected if the change in D_c_ was wholly responsible for the pre – post TSD differences in rates of state expression. This line explained 60%, 55.5% and 0% of the variance in the data in WAKE, NREM and REM, respectively, according to adjusted R^2^.

## 4. Discussion

The present study reveals an important role for ultradian rhythms in influencing the amplitude of sleep-wake circadian rhythms, the magnitude of post-TSD rebound, and in the overall rates of state expression during steady baseline. The data give new insight into the interrelations between short-term state transitions, medium-term ultradian timing mechanisms and long-term patterns of sleep-wake state in rats. Ultradian timing is strongly implicated in the control of all three states of wakefulness, NREM sleep and REM sleep, and appears to represent an important mechanistic link between short-term probabilistic state transitions and longer-term deterministic sleep-wake patterns.

### 4.1. Estimation of the quantitative influence of ultradian timing on sleep-wake patterns

Post-TSD rebounds in NREM, total sleep and WAKE were present when the data were calculated on a full ultradian cycle-by-cycle basis but absent in each of the υ_s_ and υ_w_ phases individually (Fig 8). How can there be a rebound over a full cycle if there is no rebound in either of its two phases? Also, it is well known that ultradian and circadian rhythms occur concurrently in an LD cycle (Fig 3), but how do they interact? The answers to these and other questions can be revealed by quantitative exploration of the relative importance of ultradian timing versus ultradian amplitude (i.e. phase-specific state content) on long-term sleep-wake patterns.

In this study, the rate of expression of state i was quantified as fractional time in state i (i.e. F_i_ = time in state i / total time, where subscript i denotes _WAKE, NREM_, or _REM_). Long-term patterns of behaviour that are clearly visible in the hypnogram, such as diurnal rhythms and post-TSD rebounds, reflect corresponding variation in F_i_. Long-term F_i_ can be altered in two additive ways. The first is a change in bout-to-bout state transition probabilities leading to altered durations and/or frequencies of state bouts.

Alternatively, a change in F_i_ is achievable in principle without invoking adjustments at the level of state bout transitions. This is the case if the hypnogram features oscillatory patterns (e.g. ultradian and circadian rhythms) in which each phase of an oscillation is characterized by a distinct regime of transition probabilities. In this scenario, the change in long-term F_i_ is brought about not by varying the actual transition probabilities within one or both phases, but by a change in the fraction of time per cycle that is devoted to the two different phases (the ultradian “duty cycle”, D_c_). Variation in D_c_ constitutes adjustment of an ultradian timing mechanism.

Quantification of the impact of the ultradian timing mechanism on long-term F_i_ requires knowledge of two variables; the difference in F_i_between the two ultradian phases (υ_s_-υ_w_F_i_), and the magnitude of the change in D_c_ (ΔD_c_):

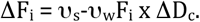

Fig 7G,H,I shows that the post-TSD rebound included a transient reduction in D_c_ over the first 3-5 ultradian cycles in LL animals. This transient was most pronounced in the SD6 group (initial ΔD_c_ = -0.2, falling to 0 over 4 cycles). Using values given in Table 1 for LL rats, we calculate that in BL, υ_s_-υ_w_F_WAKE_ ≈ -0.63, υ_s_-υ_w_F_NREM_ ≈ 0.54 and υ_s_-υ_w_F_REM_ ≈ 0.09. If peak ΔD_c_ = -0.2, to what extent does this account for the observed post-TSD rebounds over the first REC day in the SD6 group?

In the SD6 group of rats, mean WAKE excess was 234 min, and WAKE rebound was -94 min (40%), mean NREM deficit was -202 min, and NREM rebound was 69 min (34%), and mean REM deficit was -32 min, and REM rebound was 25 min (77%). Mean ΔD_c_ over 6 ultradian cycles (1 day) was -0.083. Hence, mean ΔF_WAKE_ due to ultradian timing was -0.63 x -0.083 = -0.0525, which represents a change of 0.0525 × 24 h, or 76 min. This is ∼80% of the mean ΔF_WAKE_ (−0.0653) observed in the rebound of the SD6 group. Similar analyses yield estimates of 94% for the NREM rebound and 43% for the REM rebound.

This simple calculation indicates that under the present experimental conditions, a change in ultradian timing underlies most, but not all, of the rebounds in WAKE and NREM, and less than half of the observed rebound in REM. This further implies that alterations in state transition probabilities within the ultradian cycle also play a role in setting the magnitude of the rebound, especially for REM. Fig 8 shows that the latter effect is present in the υ_s_ phase only. We confirmed previous reports of little or no correlation between WAKE bout duration and subsequent sleep episode duration[9, 23, 24]. We also found no effect of TSD on the positive correlation between REM bout duration and subsequent interREM interval, which has been interpreted as a REM homeostat [12].

Transition probabilities within the υ_w_ ultradian phase were unaffected by TSD and played no direct role in the ensuing rebound in any states. This refutes our previous prediction that the high “gain” of state maintenance probability of the “long” WAKE sub-type, which is mostly restricted to the υ_w_ ultradian phase, would be the main mechanism underlying the rebound response[37].

A similar calculation can be used to estimate the relative importance of within-phase state transition versus between-phase ultradian timing in determining the amplitude of sleep-wake diurnal rhythms under LD cycles. Our reanalysis of data from 10 rats entrained to a 12:12 h LD cycle found that both ΔD_c_ (Fig 2) and υ_s_-υ_w_F_i_ (Fig 3) differed between L and D. Using data presented in Table 1 we estimate that within-phase changes in F_i_ (i.e. state transition probability adjustments) account for approximately 72% of the diurnal rhythm amplitudes in all three states, while L to D variation in ultradian D_c_ account for the remaining 28%. This indicates that there is a direct and reciprocal quantitative link between ultradian and diurnal rhythm amplitudes under LD cycles. The circadian system influences ultradian amplitude by modulation of F_i_ values in both υ_s_ and υ_w_ (Fig 3). At the same time, ultradian timing mechanisms influence diurnal rhythm amplitude via variation of D_c_ across the LD cycle (Fig 2).

Finally, we observed that TSD was often followed by variation in steady baseline rates of sleep-wake expression (Fig 4). This effect was not correlated with TSD duration, or estimated circadian phase at REC onset. The absolute changes were small and our calculations, based on the principles described above, indicate that the magnitude of the change in rate was largely accounted for by changes in ultradian D_c_ in WAKE and NREM but not in REM (Fig 13). In contrast to WAKE and NREM, baseline rate of REM expression was not affected consistently by the TSD-induced changes in D_c_.

### 4.2. Implications for study design and analysis

An issue that comes into sharp focus in the present study concerns sources of sample heterogeneity that can shape overall responses. An observed change in a sample mean quantity such as bout duration is conventionally assumed (usually implicitly) to reflect direct control via state-switching neural mechanisms[52]. However, this interpretation is predicated on the assumption that the dataset is homogeneous. When this is not the case, such as when ultradian and circadian oscillations are present, then an indirect “phase blending” phenomenon can lead to variation in estimated mean bout durations and transition probabilities without the direct involvement of such neural mechanisms.

This phenomenon has practical implications for the design of data sampling protocols in studies of sleep-wake state (and indeed any other biological variables that display ultradian rhythms). Since ultradian cycles are quasi-periodic and often asymmetric (i.e. D_c_ ≠ 0.5), standard approaches of using fixed sampling windows risk incurring errors that can be avoided by a sampling protocol that explicitly accounts for ultradian phase. To give a simple hypothetical example of the potential importance of this issue, imagine a study with the objective to determine whether there are changes in F_NREM_ between the first (ZT0-6) and second (ZT 6-12) halves of the L phase of the diurnal cycle. We use the data in Table 1, with some simplifying assumptions: assume constant F_NREM_ across the entire L phase = 0.55, constant D_c_ = 0.34, constant F_NREM_ in υ_w_ = 0.33, constant F_NREM_ in υ_s_ = 0.66, constant ultradian period = 4 h and synchronous L phase and υ_s_ phase onsets. Taking a simple average across each ZT0-6 and ZT6-12 interval would yield F_NREM_ ZT0-6 = 0.59, F_NREM_ ZT6-12 = 0.51. This yields the erroneous conclusion that the NREM content differs between early and late L phase, with both values of F_NREM_ differing from the known correct (and constant) value of 0.55. This error arises because of an asymmetry in the distribution of υ_s_ and υ_w_ phases in the two halves of the L phase. ZT 0-6 consists of 4.64 h in the υ_s_ phase and 1.36 h in the υ_w_ phase, whereas ZT6-12 consists of 3.28 h in the υ_s_ phase and 2.72 h in the υ_w_ phase. Other quantities such as bout durations, bout frequencies, transition probabilities etc., if they differ significantly between υ_s_ and υ_w_ phases, would be affected in the same way by the unequal mix of ultradian phases in the two measurement intervals.

This “back-of-the-envelope” calculation serves to highlight the potential pitfalls of failing to quantify and account for ultradian rhythms during data analysis. Exactly the same problem would arise should we fail to account for circadian phase in data sampling. However, unlike circadian rhythms, ultradian oscillations are quasi-periodic and so individual phase durations cannot be predicted *a priori*. It is not safe to assume that a 2 h window of time at a given circadian phase will contain the same proportions of υ_s_ and υ_w_ on two consecutive days. Hence, the ultradian phases must be measured directly for accurate analysis.

### 4.3. Ultradian and circadian rhythms in LL

This study has confirmed previous reports of profound attenuation of circadian rhythm amplitude with ongoing ultradian rhythmicity in rats kept in prolonged bright light (LL) [53-55]. Continuous wavelet analysis provided the first clues of a potential link between ultradian rhythms and the expression of circadian rhythms and post-TSD rebounds, demonstrating an important advantage of this method[41] for analysis of non-stationary rhythms over time. Fig 1 panels A-C illustrate a complete 25-day study in a representative rat in LL. The vertical grey bars denote ∼4 h TSD and the spectra show a subsequent transient perturbation of power in the ultradian period range (2 – 8 h), suggesting post-TSD change in ultradian characteristics. Power in the circadian period range was weak and ephemeral, with a generally stronger signal in T_b_ than F_WAKE_. Panel C serves to confirm the visual impression from panels A and B that T_b_ and F_WAKE_ co-varied over time. Similar to mouse T_b_[56], the peak periods of ultradian oscillations drifted considerably over time and there were sometimes two or more concurrent periods within the same part of the time series. For comparison, panel D illustrates a week of baseline F_WAKE_ under an LD cycle. The strong wavelet power at 24 h is a prominent feature of this spectrum, which contrasts strikingly with the LL rat (B). Also noteworthy are the non-steady ultradian periodicities, which bear similarities to the LL condition, but with one important difference; in LD there was a clear diurnal modulation of the signal with suppression of power within the 2-3 h period range over the L phase of each day (plotted as the first half of each day on the time axis).

### 4.4. Sleep-wake homeostasis and the magnitude of post-TSD rebounds in LL

This study demonstrates that a monotonic post-TSD rebound response can occur under conditions of constant light and severely suppressed diurnal rhythmicity. However the rebound responses were highly variable, both within and between animals, and apparently absent in about a third of all trials (Fig 4). Furthermore, in those trials that did exhibit a monotonic rebound, the responses of WAKE and NREM came to completion long before full compensation had been realized, at around 50%. This result is in broad agreement with previous studies of rats whose circadian rhythms were abolished by surgical lesions of the suprachiasmatic nuclei [57, 58]. Mistlberger et al [57] reported data for 3 days following 24 h TSD in rats with and without circadian rhythms, and in the presence and absence of LD cycles in lesioned animals. They reported little difference between conditions; rebounds were incomplete, at approximately 16 – 26% for t_NREM_ and approximately 29 – 34% for t_REM_. Collectively, the results of these studies suggest that circadian rhythms contribute to post-TSD rebounds, further supporting suggestions that the two processes are not independent[59].

By contrast, in the present study the group mean rebounds were essentially complete for REM sleep (Fig 6), also corroborating several previous reports for rats in an LD cycle[5, 17, 47, 60]. This suggests that the REM response differs to some degree from that of NREM in terms of its underlying mechanism. In the present study we find that the REM rebound depends to a greater extent on changes in NREM transition trajectory within the υ_s_ ultradian phase (Fig 11).

We observed that ultradian rhythms continue unabated during post-TSD rebound. Moreover, there was no consistent prolonged extension of the υ_s_ phase at the onset of rebound, contradicting the prediction of our simple model of quasi-periodic ultradian rhythm[19]. In fact, we observed the opposite – a response mainly mediated by a shortening of the υ_w_ phase of the ultradian cycle (Fig 7). Accordingly, we conclude that the model is implausible and reject it as a useful framework for furthering our understanding of ultradian rhythm generation.

A concept that has been proposed to explain the disconnect between t_NREM_ deficit and ensuing t_NREM_ rebound, is that an “intensity” dimension contributes to NREM sleep expression[50], and that intensity is correlated with slow waves in the EEG during NREM (SWA_NREM_). There is strong evidence that high SWA_NREM_ stages of NREM are preferentially maintained under conditions of sleep restriction and compensated following their suppression[61-63]. Agnew and colleagues suggested that SWA_NREM_ (stage 4 sleep) is a functional necessity[64-66] and Feinberg[51] proposed that SWA_NREM_ is regulated by a control mechanism that serves to maintain WAKE-related brain metabolic function (the delta homeostasis hypothesis). Borbély incorporated this hypothesis into his early descriptive version of the two-process model of sleep regulation[50]. Feinberg’s model predicts that a TSD-induced deficit in cumulative SWA_NREM_ (SWA_NREM_ x t_NREM_ = SWE_NREM_) should be compensated during recovery. However, full compensation was not confirmed in this (Fig 12) nor in other studies[10, 60, 67].

Our present data corroborate previous reports[5, 10, 47, 68] of a biphasic response in SWA_NREM_ following TSD in rats; an initial positive rebound, which in the present study was found to occur only in the ultradian υ_s_ phase, followed by a negative rebound in both ultradian phases (Fig 12A,B,C). Combining this with t_NREM_ (Fig 8D,E,F), a monotonic rebound in SWE_NREM_ was observed in υ_s_ only, but it incompletely compensated for the TSD-induced SWE_NREM_ deficits (Fig 12G,H,I). Indeed, the rebound in SWE_NREM_ was no more complete than the rebound in t_NREM_ alone, owing to the fact that the negative rebound in SWA_NREM_ cancelled the preceding positive rebound yielding no net contribution from SWA_NREM_ to the final magnitude of the rebound in SWE_NREM_. This conclusion holds in relation to each ultradian phase and over full ultradian cycles. We conclude from this that these data are inconsistent with the hypothesis that SWE_NREM_ is homeostatically regulated, and support a growing consensus that SWA_NREM_ is not a direct or complete quantitative index of NREM “intensity”[59, 69-72].

### 4.5. Behavioural homeostasis versus homeostatic behaviour

The concept of sleep homeostasis has been a central pillar of theories on the regulation and function of sleep since it was introduced by Borbély[73]. Although the usage of this term is sometimes ambiguous, in the present paper we follow Borbély’s definition: “The term ‘sleep homeostasis’ posits that sleep strives to maintain a constant level by variation of its duration and intensity”[74], with “the latter being reflected by SWA”[8]. This hypothesis represents a model of *behavioural homeostasis*, in which NREM sleep is the regulated variable, and time in state (t_NREM_) and state “intensity” (indexed by SWA_NREM_) are the putative effector variables. Despite widespread acceptance over several decades within the field of sleep research[75], the general validity of the notion that behaviour per se is regulated by homeostatic control mechanisms has long been a subject of debate[76, 77] and has not yet been satisfactorily resolved. In the case of sleep homeostasis, it is possible to rationalize partly incomplete rebounds in terms of supplemental hypotheses such as “core sleep”[78, 79] or nonlinear error signals[80], but such ideas, while interesting, are highly speculative and in the absence of a reliable metric for homeostatic error signal, are currently untestable. As mentioned earlier, SWA_NREM_ is proving to be increasingly inadequate for that purpose[59, 69-72]. We entertained some of these supplemental ideas in our previous study, where shortfalls in rebound compensation were not too great[10], and concluded that we could not rule out a mechanism of behavioural homeostasis. However, in the present study the discrepancies between observation and prediction would seem to be too extreme to be accounted for by such modifications of the basic homeostatic proportional control model. It is possible that the experimental conditions of LL had a disruptive effect on the control system, but that too is a matter of conjecture at the present time.

The model of sleep as a homeostatically controlled variable is distinct from a model of sleep as an effector mechanism in the homeostatic control of some other functional system (i.e. a model of “*homeostatic behaviour*”). Numerous examples of homeostatic behaviour have been documented, including feeding behaviours, thermoregulatory behaviours (e.g. huddling)[77] and respiratory behaviours (e.g. resurfacing in marine mammals[81]). Beersma and Daan[80] have added sleep to this list, claiming that “Sleep is part of the processes contributing to brain homeostasis; it is not the regulated variable itself” and “there is no a priori reason to expect that sleep itself, whether measured in minutes or in energy (integrated slow wave activity in the EEG), should be homeostatically defended.” The data presented here lend support to this assertion.

The model of sleep as a homeostatic behaviour postulates that sleep is an “on-demand” effector response involved in the homeostatic regulation of putative state-dependent functions. It is likely that a wide range of central neural (and bodily) functions are associated with sleep, not all of which will necessarily be homeostatic, and even those that are may not necessarily coincide in their demands for sleep. To refer to this kind of control as “sleep homeostasis” is therefore misleading and inapproriate.

The above discussion suggests that perhaps it is time to look outside of the conceptual confines of prevailing dogma and search for alternative explanations for observed phenomena. Both of the above “sleep homeostasis” models (behavioural homeostasis and homeostatic behaviour) make the fundamental assumption that sleep is the target of the control system. However this is not necessarily the case; the results of this and other studies can also be rationalized in terms of control of wakefulness[10, 37]. Aside from the above-mentioned fact that there are many known homeostatic functions that depend upon wake-related behaviours (whereas in contrast unequivocal sleep-dependent homeostatic functions have proven more elusive), evidence is rapidly accumulating that WAKE-dependent motivational, emotional and other behavioural factors have a profound impact on overall patterns of state expression[13, 82-85]. On the basis of the results in the present study, we hypothesize that some effects on vigilance state of such factors as environmental temperature[86], stress[87], energy balance[83] and social interactions[88] might be mediated, at least in part, by adjustments in ultradian phase and amplitude.

### 4.6. Critique

In this study, the application of constant bright light to suppress circadian rhythms was not entirely successful in suppressing the circadian rhythm. Furthermore, the very low and variable amplitude of the residual free-running circadian rhythm, as shown in wavelet spectra (Fig 1), led to some uncertainty regarding the assignment of circadian phase in some animals. We were therefore unable reliably to ascertain whether TSD caused circadian phase shifts and, if so, whether they were linked to changes in ultradian phase. This highlights the need for a more comprehensive study of animals in constant darkness (DD) to examine the interactions between post-TSD rebound, ultradian and circadian rhythmicity.

Prolonged LL led to suppression of circadian variations in ultradian parameters; phase durations, duty cycle (Fig 2), amplitude and phase-specific fractional state content (Fig 3). As explained earlier, these factors combine to determine overall state expression, which in LL exhibited lower F_WAKE_ and F_REM_ (by 9.2 % and 30.8 % respectively) and higher F_NREM_ (by 19.3 %) compared with LD in this study. The extent to which these changes are attributable to suppression of circadian amplitude versus direct masking affects of light cannot be determined from this study design and further research is needed to address this question.

In this study we divided the rats into 3 groups of animals to test the hypothesis that magnitude of post-TSD rebound is proportional to TSD-induced sleep-wake perturbation. The disadvantage of this was that each group had relatively small sample sizes, a problem that was exacerbated by loss of an animal from each group due to equipment malfunction, the unanticipated failure of many trials to exhibit “normal” monotonic rebounds, and the above-mentioned failure to completely suppress circadian rhythmicity. This issue was particularly acute in analysis of the υ_w_ phase, which had fewer state transitions than the υ_s_ phase. All of these problems inevitably compromised the statistical power of the analyses. Future work must include larger sample sizes for more reliable statistical tests.

Given that the basic aim of the study was to examine the interaction between ultradian rhythms and post-TSD rebounds, we felt that inclusion of animals that displayed no apparent rebound might conceal important features and possibly lead to erroneous conclusions. However, the decision to exclude these trials clearly relied upon a preconceived notion of how the response should play out, based upon classical concepts of sleep homeostasis. If, as we have suggested, the post-TSD rebound is a manifestation of altered WAKE-related motivations, then the absence of a visibly nonlinear rebound would not necessarily be a valid reason to exclude the data from analyses. At this stage, this issue remains a matter of conjecture and further research targeting motivational drives and ultradian regulation will be needed to clarify these uncertainties.

### 4.7. Concluding remarks

In the present study we show that long-term baseline rates of state expression (Fig 13), circadian rhythm amplitude (Figs 2,3) and post-TSD rebound magnitude (Figs 5,6) are all closely linked to ultradian rhythm expression in rats. Two main mechanisms were identified as important determinants of rebound magnitude: ultradian phase-specific transition probabilities, and duration of the WAKE-dominant υ_w_ phase of the ultradian cycle. Of the former, rebounds were enhanced by decreased probabilities of arousal from NREM 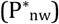 in the υ_s_ phase (Fig 11). This ensemble of WAKE-related responses is consistent with a hypothesis that the post-TSD rebound results from a modification of WAKE-related drives[10, 37]. Regulation of the timing of the WAKE-dominant υ_w_ phase of the ultradian cycle provides an important mechanism by which long-term patterns can emerge in both wakefulness and sleep, without direct modulation of the neural systems mediating stochastic state transitions.

This study has emphasized that the ultradian rhythm is an important source of sample heterogeneity, which should be accounted for in analyses. Moreover, the episodic[56] or “jaggy”[89] appearance of ultradian rhythms may be less random than it seems, and at least some of the variability might instead reflect ongoing controlled adjustment of phase and amplitude to mediate patterns of expression of all three sleep-wake states.

## Acknowledgements

This research was supported by a Discovery Grant from the Natural Sciences and Engineering Research Council of Canada.

